# NLRP3 is a thermosensor that is negatively regulated by high temperature

**DOI:** 10.1101/2025.09.29.679254

**Authors:** Wei Wang, Damien Bertheloot, Junya Zhang, Marcia A. Munoz, Shangze Xu, Amelia Stennett, Chloe M. McKee, Ryan Knight, Emma C. McKay, Bernardo S. Franklin, Michael J. Rogers, Agnieszka K. Bronowska, Rebecca C. Coll

## Abstract

Inflammation is an essential response to infection and injury, but unregulated inflammation is damaging and must be limited by negative feedback signalling. Inflammasome signalling drives local inflammation and systemic responses like fever. However, our understanding of how inflammasome signalling is negatively regulated is limited. NLRP3 is activated by a vast number of stimuli and senses perturbations of cytoplasmic homeostasis. As temperature is a fundamental environmental stressor, we hypothesised that NLRP3 inflammasome signalling would be sensitive to increased temperatures and so we investigated the effects of high temperatures on NLRP3 in macrophages. Short-term incubation at high fever range temperatures significantly inhibits NLRP3 activation, while secretion of the inflammasome-independent cytokines TNF and IL-6 are much less affected. High temperature blocks NLRP3 inflammasome formation in a transcription-independent manner, and NLRP3 is highly sensitive to temperature-mediated inhibition relative to the NLRC4, AIM2, and NLRP1 inflammasomes. Using cellular assays and molecular simulations we show that the effect of high temperature on NLRP3 is protein intrinsic. NLRP3 activation is associated with a decrease in the thermal stability of the protein and multiscale molecular dynamics simulations identified a peptide in the <u>C</u>-terminal <u>o</u>f the <u>FI</u>SNA domain (COFI) that is highly flexible and undergoes a significant conformational shift at high temperature. Cellular assays demonstrate that the COFI regulates NLRP3 stability and is required for activation. Furthermore, mice exposed to high temperature display attenuated inflammatory cytokine production upon *in vivo* LPS challenge. Our studies thus reveal that high temperatures associated with fever limit NLRP3 activity and identify a novel role for NLRP3 as a protein thermosensor.

## Introduction

Inflammation is an essential host response that can be locally restricted, or in the case of disseminating infections or severe injury can trigger systemic responses such as fever^2^. Fever is an ancient and highly conserved element of the immune response, and calor or heat was recognised by Celsus in the first century BC as one of the cardinal signs of inflammation^3^. Fever range temperatures (FRT) can have direct effects on pathogen replication but also regulate the immune response^3,4^. FRT enhances both the recruitment of neutrophils to sites of infection and the respiratory burst, while it increases lymphocyte recruitment and the phagocytic capacity of macrophages^3,5^. More recent work demonstrates that FRT also boosts the differentiation and pathogenicity of Th17 and Th1 cells^6,7^, highlighting the important effects of temperature on the immune system.

Unregulated inflammation or sustained fever is damaging and must be limited by negative feedback signalling^3^. Inflammation and fever are initiated via the production of inflammatory cytokines such as interleukin (IL)-6, TNF, and IL-1β produced by cells of the innate immune system in response to pathogen- and damage-associated molecular patterns (PAMPS/DAMPS)^2,3^. IL-1β is the most potent cytokine inducer of fever, and its production is tightly regulated^10^. Inflammasomes are intracellular protein complexes activated by PAMPS and DAMPs which orchestrate the activation of inflammatory caspases. Caspase-1 cleaves the inflammatory cytokines pro-interleukin(IL)-1β and pro-IL-18 into their active secreted forms, and cleaves Gasdermin-D (GSDMD) leading to pyroptosis^8,9^. Inflammasome formation is initiated by sensor proteins including members of Nod-like receptor (NLR) family such as NLR family pyrin domain containing 3 (NLRP3) and inflammasome signalling is a highly inflammatory process.

NLRP3 activation can be triggered by a huge array of stimuli, varying from pore forming toxins and crystals to ionic fluxes and disruption of endosomal trafficking^11,12^. NLRP3 is therefore regarded as a general sensor of disturbance in cellular homeostasis^13,14^. Autosomal dominant mutations in NLRP3 cause a group of autoinflammatory diseases (AIDs) called cryopyrin-associated periodic syndromes (CAPS). In one of these rare conditions, familial cold autoinflammatory syndrome (FCAS), NLRP3 is activated by cold temperatures. Individuals with FCAS experience symptoms such as urticaria-like rash and fever following exposure to cold^14^. In addition to NLRP3-AIDs, NLRP3-dependent inflammation is associated with the pathogenesis of many chronic human diseases including Parkinson’s and atherosclerosis. NLRP3 inhibition is therefore a promising therapeutic strategy that is currently in clinical trials^15,16^. Despite significant advances in our understanding of NLRP3 activation and the development of specific inhibitors, negative regulation of NLRP3 is less well characterised.

Prior to the description of the inflammasome, several studies examined the effects of temperature on cytokines and identified that IL-1β production was regulated by FRT^17–20^. More recent preliminary work linked high temperatures to the inhibition of inflammasomes, but the underlying mechanisms are unknown^21–23^. Recent studies observed that macrophages subjected to heat stress (43°C) undergo a necrotic cell death programme dependent on receptor-interacting protein kinase 3 and ninjurin-1^24,25^. In addition, heat stress at 43°C was shown to activate NLRP3-dependent caspase-1 cleavage^25^. Given the association of NLRP3 with cold temperatures and its role in driving the production of fever-inducing cytokines, we hypothesised that NLRP3 would be sensitive to FRT.

Here we describe how high temperatures affect NLRP3 activation. We find that short exposure to high temperatures (39.5-42°C) inhibits NLRP3 activation in mouse and human macrophages. NLRP3 is highly sensitive to increased temperatures and its activity is regulated independent of transcription. Using complimentary *in silico* and *in cellulo* analyses we find that NLRP3 is regulated by temperature in a protein intrinsic manner. We identify a specific peptide in the NLRP3 FISNA domain that senses temperature and is required for NLRP3 inflammasome formation. We also use an *in vivo* approach to examine the effect of temperature on inflammatory responses and observe that FRT negatively regulates lipopolysaccharide (LPS)-induced inflammation in mice. Finally, we show that high temperature attenuates the formation of ASC specks in cells expressing hyperactive NLRP3 mutations associated with CAPS, pointing to the therapeutic relevance of this process. We have identified a novel endogenous feedback mechanism that limits damaging NLRP3-associated inflammation and propose that NLRP3 functions as a protein thermosensor.

## Results

To model the effects of high temperature in cells expressing high levels of NLRP3^26^, we used mouse bone marrow-derived macrophages (BMDM) primed with LPS at 37°C for 3 hours (h). BMDM were then incubated at 42°C for 1 h before being returned to 37°C (Supp. Fig. 1A). Heat shock at 42°C did not trigger inflammasome activation in LPS-primed cells as there was no active IL-1β release, minimal pyroptosis measured by lactate dehydrogenase (LDH) release, and no caspase-1 or IL-1β cleavage (Fig. 1A-C). However, incubation at 42°C prior to the addition of the potassium ionophore nigericin to activate NLRP3 significantly limited IL-1β, pyroptosis, and both caspase-1 and IL-β cleavage (Fig. 1A-C). The heated BMDM were still capable of cytokine secretion as the production of TNF production was only slightly affected (Supp. Fig. 1B). Similar to the effect of high temperature, the specific NLRP3 inhibitor MCC950^27^ blocked inflammasome responses but not TNF release at 37°C (Fig. 1A-C, Supp. Fig. 1B).

**Figure 1.**
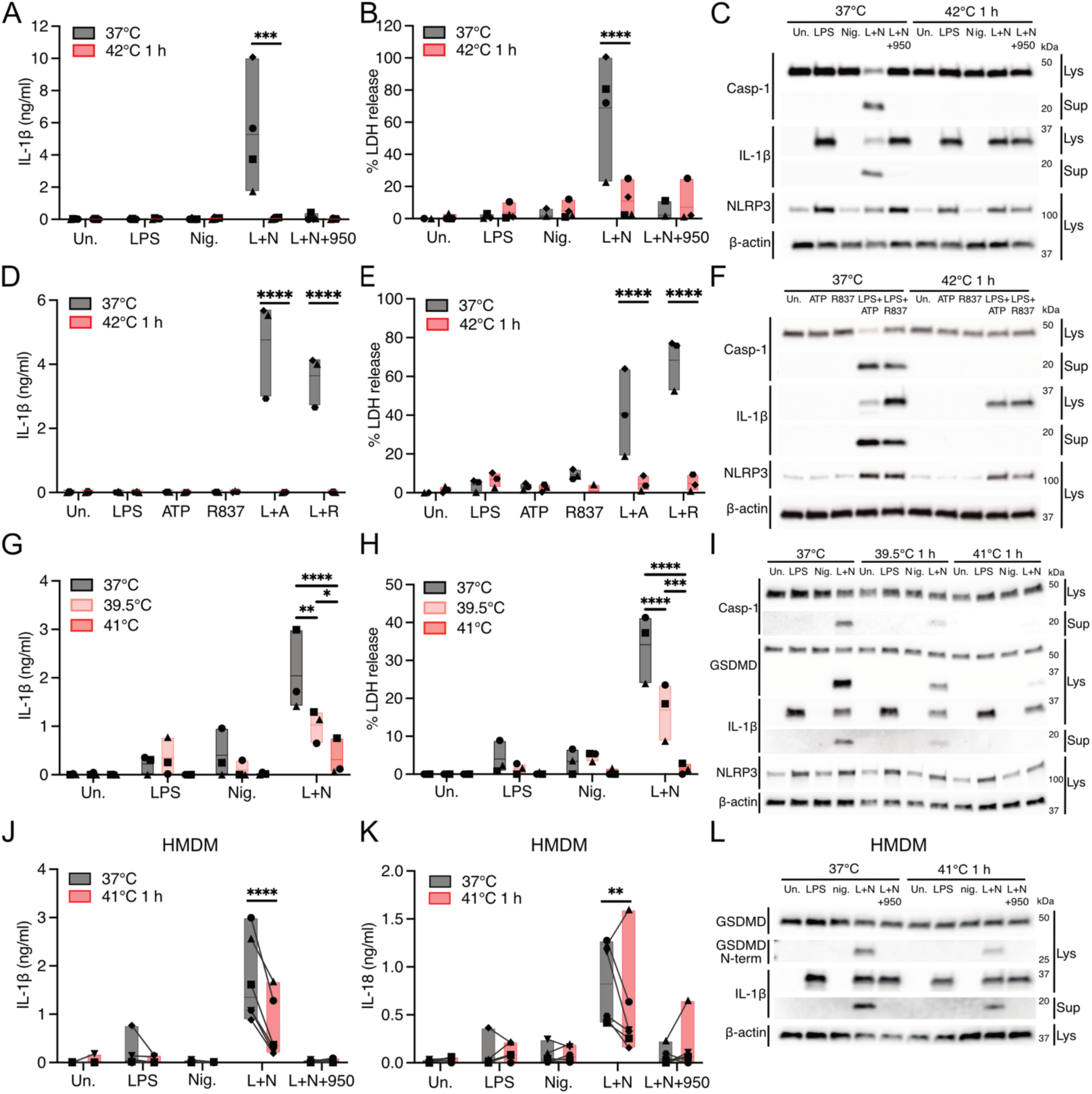
NLRP3 inflammasome signalling is inhibited by short term incubation at high temperatures. (A-F) BMDM primed with LPS (100 ng/ml) for 3 h at 37°C. MCC950 (100 nM) or H2O vehicle control was then added before half the cells were moved to 42°C for 1 h. All cells were then stimulated with nigericin (5 μM), ATP (2 mM), or R837 (100 μM) at 37°C for 1 h. (G-I) iBMDM were primed with LPS (100 ng/ml) for 3 h at 37°C before cells were moved to 39.5 °C or 41°C for 1 h. All cells were then stimulated with nigericin (5 μM) at 37°C for 1 h. (J-L) HMDM were primed with LPS (100 ng/ml) for 3 h at 37°C. MCC950 (100 nM) or H2O vehicle control was then added before half the cells were moved to 41°C for 1 h. All cells were then stimulated with nigericin (5 μM) at 37°C for 2 h. Cell free supernatants were analysed by ELISA for IL-1β (A, D, G, J) and IL-18 (K), and by LDH release assay (B, E, H). Supernatants and cell lysates were analysed by western blotting for cleavage and expression of Caspase-1, IL-1β, GSDMD, and NLRP3. β-actin is used as a loading control (C,F,I,L). Average value of three technical replicates from N=3-4 independent biological replicates (symbols), floating bars (min to max) line at mean (A, B, D, E, G, H). Average of three technical replicates from N=6 donors indicated by symbols linked by lines, floating bars (min to max) line at median (J, K). Representative blots from N=2-4 biological replicates (C, F, I, L). Data were analysed by Two-way RM ANOVA with Šídák’s multiple comparisons test p= <0.05* p< <0.01**; < 0.001***.< 0.0001****.

**Supplemental Figure 1.**
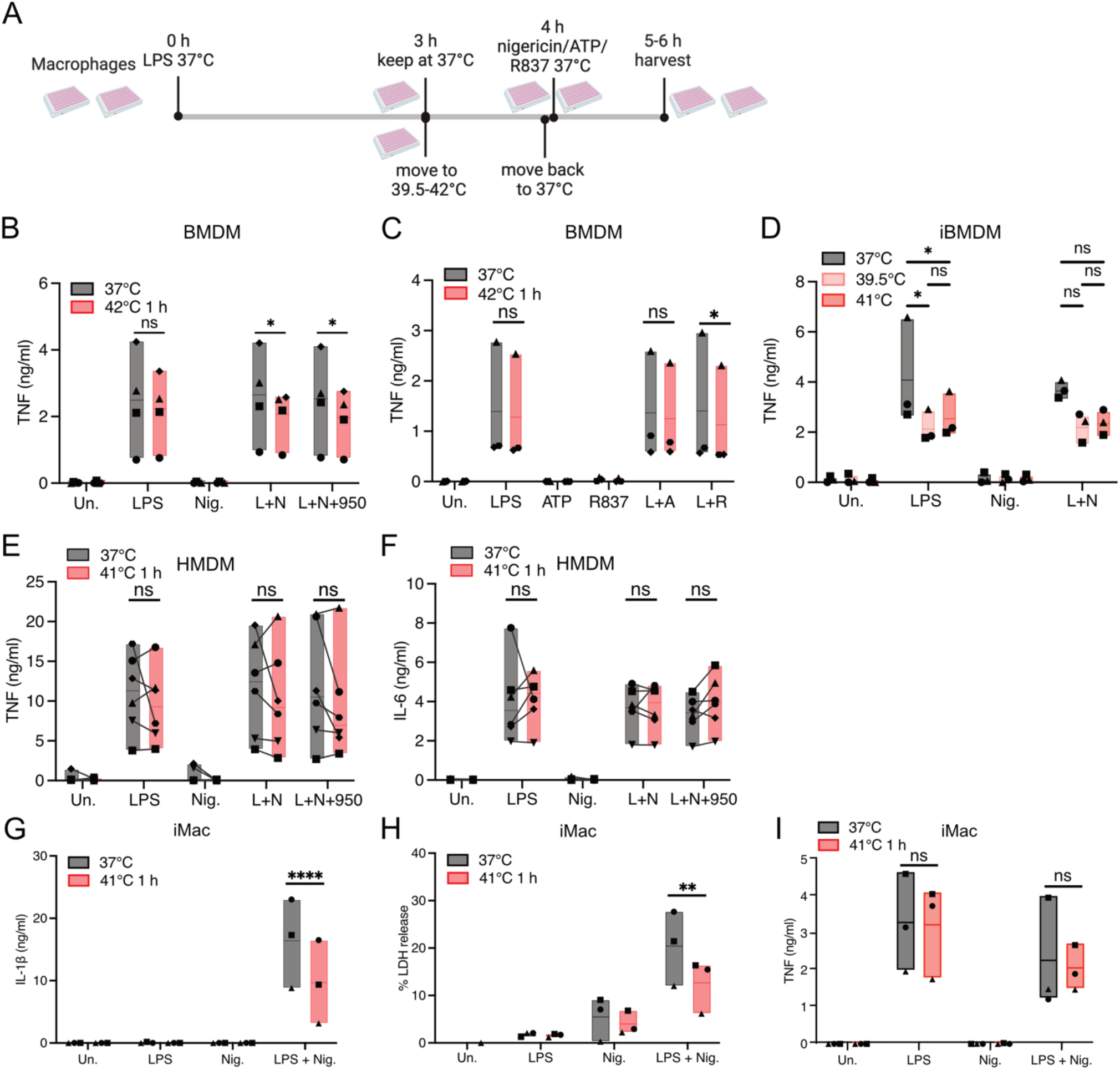
N**L**RP3 **inflammasome signalling is inhibited by short term incubation at high temperatures.** (A) Schematic diagram of the experimental approach used for Figure 1 and Supp. Fig 1. (B-C) BMDM primed with LPS (100 ng/ml) for 3 h at 37°C. MCC950 (100 nM) or H2O vehicle control was then added before half the cells were moved to 42°C for 1 h. All cells were then stimulated with nigericin (5 μM), ATP (2 mM), or R837 (100 μM) at 37°C for 1 h. (D) iBMDM were primed with LPS (100 ng/ml) for 3 h at 37°C before cells were moved to 39.5 °C or 41°C for 1 h. All cells were then stimulated with nigericin (5 μM) at 37°C for 1 h. (E-I) HMDM or iMacs were primed with LPS (100 ng/ml) for 3-4 h at 37°C. MCC950 (100 nM) or H2O vehicle control was added where indicated, before half the cells were moved to 41°C for 1 h. All cells were then stimulated with nigericin (5 μM) at 37°C for 2 h. Cell free supernatants were analysed by ELISA for TNF (B-E, I), IL-6 (F), IL-1β (G), and by LDH release assay (H). Average value of three technical replicates from N=3-4 independent biological replicates (symbols), floating bars (min to max) line at mean (B, C, D, G, H, I). Average of three technical replicates from N=6 donors indicated by symbols linked by lines, floating bars (min to max) line at median (E,F). Data were analysed by Two-way RM ANOVA with Šídák’s multiple comparisons test p= <0.05* p< <0.01**; < 0.0001****.

NLRP3 activation triggered by the DAMP ATP or the potassium efflux-independent stimulus R837^21^ was also inhibited by high temperature as IL-1β secretion, pyroptosis, and caspase-1 and IL-βcleavage (Fig. 1D-F), were all blocked in heat treated BMDM, but TNF secretion was not (Supp. Fig. 1C). These results show that both potassium efflux-dependent and -independent NLRP3 activation by multiple stimuli are inhibited by high temperature. We tested the effect of lower FRT on NLRP3 activation in immortalised BMDM (iBMDM). Incubation at 39.5 °C or 41°C for 1 h also significantly decreased IL-1β release, pyroptosis, and caspase-1 and IL-1β cleavage, with minor effects on TNF release (Fig. 1 G-I, Supp. Fig. 1D). Importantly, in the heated iBMDM, pro-IL-1β and NLRP3 protein levels remained similar to those kept at 37°C (Fig. 1C, 1F, 1I). Together these data indicate that high temperatures induce a blockade of the NLRP3 inflammasome upstream of caspase-1 and IL-1β.

We next used primary human monocyte-derived macrophages (HMDM) to examine the effects of high temperature on human NLRP3. To simulated the upper limit of the normal febrile range in humans^2^, we exposed HMDM to 41°C for 1 h prior to NLRP3 activation (Supp Fig 1A). As shown in Fig. 1J, heated HMDM secreted significantly less IL-1β in response to LPS and nigericin stimulation across all donors. NLRP3-dependent IL-18 secretion was attenuated and both IL-1β and GSDMD processing were decreased in heat exposed HMDM (Fig. 1 K-L). However, levels of TNF and IL-6 (Supp. Fig. 1E-F) were not significantly decreased. We further confirmed the effects of high temperature using human induced pluripotent stem cell (iPSC)-derived macrophages (iMacs). In iMacs NLRP3-dependent IL-1β secretion and pyroptosis were significantly decreased at 41°C but TNF secretion was not (Supp. Fig. 1G-I). These data confirm that FRT negatively regulates human NLRP3 in conditions that do not block the inflammasome independent cytokines TNF and IL-6.

NLRP3 activation is tightly regulated by transcriptional priming and post-translational modifications (PTMs), such as de-phosphorylation^8^. Transcriptional priming takes hours, but NLRP3 can also be rapidly activated independently of transcription when priming and activation stimuli are added simultaneously^28–32^. We thus examined the effects of high temperature on NLRP3 activation in transcription independent conditions. NLRP3-dependent cell death was triggered by co-stimulation with LPS and nigericin in BMDM. However, cells pre-treated at 42°C did not undergo pyroptosis and caspase-1 activation was inhibited (Supp. Fig. 2A-B). In co-stimulated HMDM, pre-treatment of cells at 41°C also decreased pyroptosis and GSDMD cleavage (Supp Fig. 2C-D). Notably, pro-IL-18 is constitutively expressed in HMDM, and we observed decreased IL-18 secretion from HMDM pre-treated at 41°C (Supp Fig. 2E). High temperatures thus inhibit NLRP3 activation independent of transcriptional priming. We also tested whether high temperature inhibition of NLRP3 could be reversed upon re-exposure of cells to physiological temperature at 37°C. Interestingly, after 41°C incubation for 1 h, recovery for 2-4 h at 37°C did not rescue inhibition of NLRP3 (Supp. Fig. 2H). This shows that the effect on NLRP3 is not transient and persists for at least 4 h.

**Figure 2.**
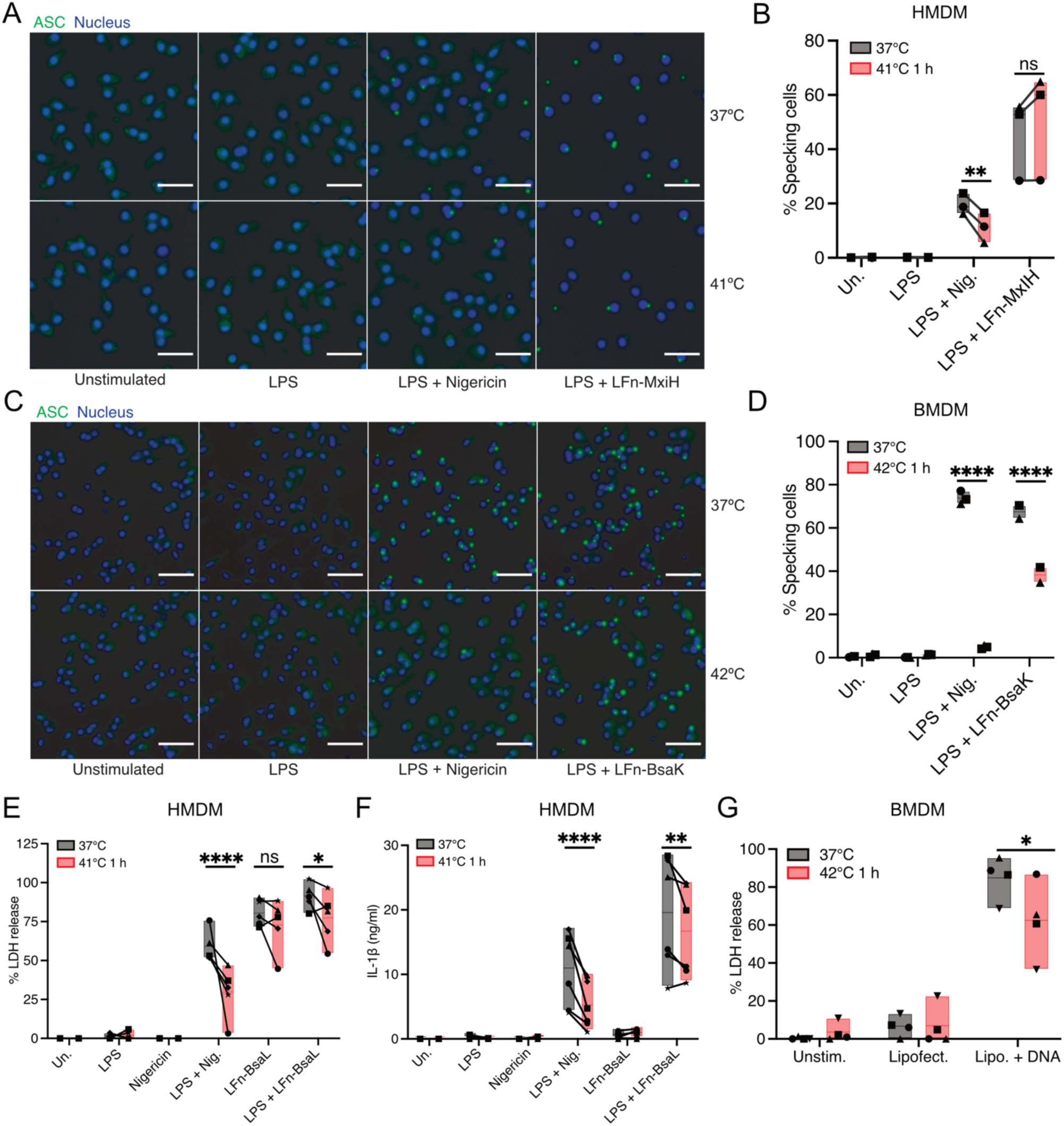
NLRP3 inflammasome formation is specifically inhibited by high temperature. (A-B) HMDM primed with LPS (100 ng/ml) for 3 h at 37°C in the presence of VX-765 (50 µM), before half the cells were moved to 41°C for 1 h. All cells were then stimulated for 2 h at 37°C with nigericin (5 μM), or LFn-MxiH (100 ng/ml) combined with PA (1 µg/ml). (C-D) ASC-mCitrine BMDM primed with LPS (100 ng/ml) for 3 h at 37°C in the presence of VX-765 (50 µM), before half the cells were moved to 42°C for 1 h. All cells were then stimulated for 2 h at 37°C with nigericin (5 μM), or LFn-BsaK (100 ng/ml) combined with PA (1 µg/ml). (A-D) Cells were fixed and stained with PE-labelled anti-ASC antibody and DRAQ5, imaged, and specks per field quantified. (E-F) HMDM primed with LPS (100 ng/ml) for 3 h at 37°C before half the cells were moved to 41°C for 1 h. All cells were then stimulated for 2 h at 37°C with nigericin (5 μM), or LFn-BsaL (1 ng/ml) combined with PA (250 ng/ml). (G) Primary BMDM kept at 37°C or 42°C for 1 h were then transfected with HT DNA (1 μg/ml) using Lipofectamine 2000 for 2 h. Cell free supernatants were analysed by LDH release assay (E, G) or by ELISA for IL-1β (F). Average values of from N=3 independent donors or biological replicates (symbols), floating bars (min to max) line at mean (B, D). Average of three technical replicates from N=5 donors or N=4 biological replicates indicated by symbols, floating bars (min to max) line at median or mean (E,F,G). Data were analysed by Two-way RM ANOVA with Šídák’s multiple comparisons test p= <0.05* p< <0.01**; < 0.0001****.

**Supplemental Figure 2.**
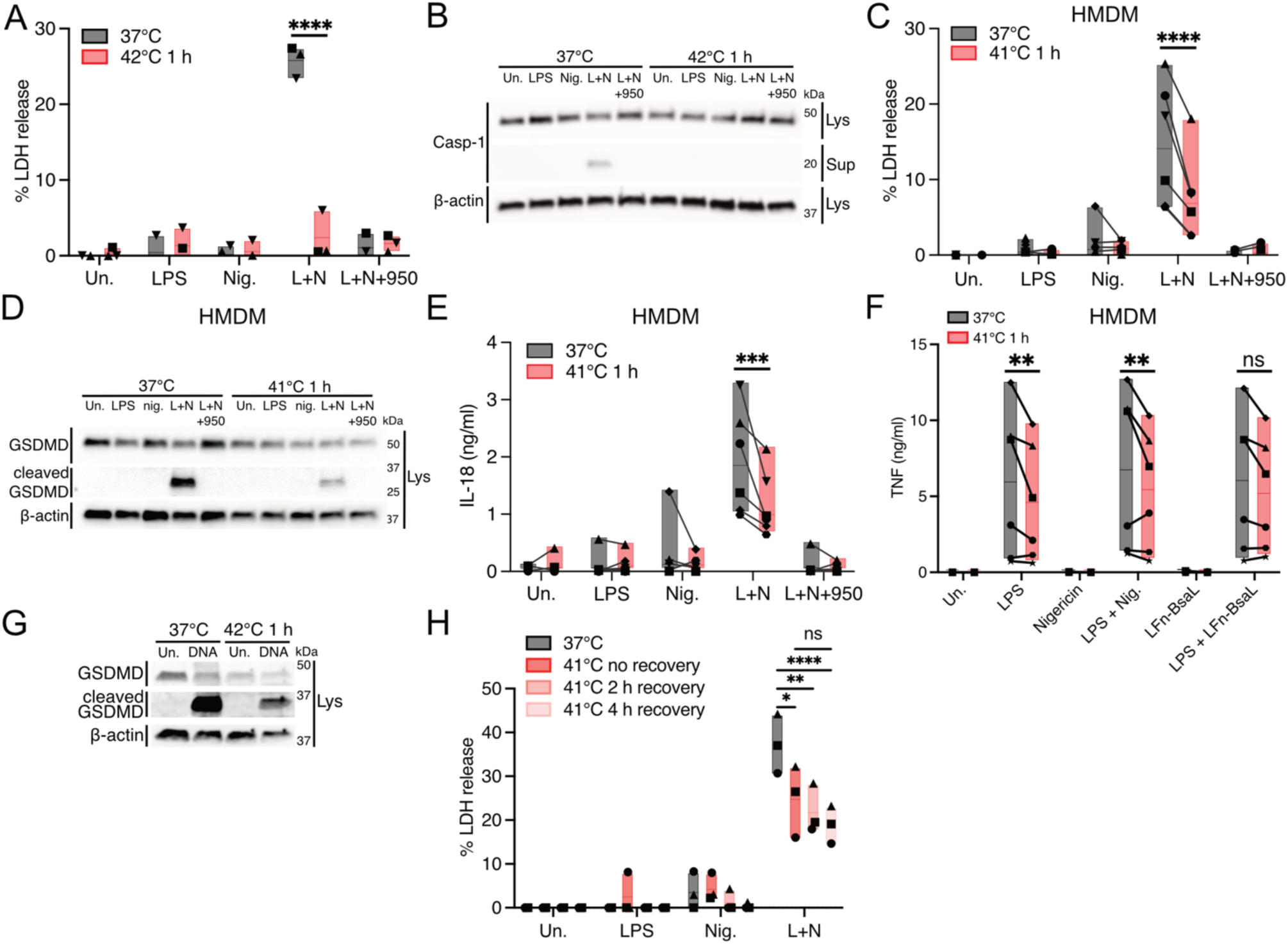
N**L**RP3 **inflammasome activation is inhibited by high temperature independent of transcriptional priming.** (A-B) Primary BMDM incubated at 37°C or at 42°C for 1 h in the presence of MCC950 (100 nM) where indicated, were then simultaneously stimulated with LPS (100 ng/ml) and nigericin (5 μM) at 37°C for 1 h. (C-E) HMDM incubated at 37°C or at 41°C for 1 h in the presence of MCC950 (100 nM) where indicated, were then simultaneously stimulated with LPS (100 ng/ml) and nigericin (5 μM) at at 37°C for 2 h. (F) HMDM primed with LPS (100 ng/ml) for 3 h at 37°C before half the cells were moved to 41°C for 1 h. All cells were then stimulated for 2 h at 37°C with nigericin (5 μM), or LFn-BsaL (1 ng/ml) combined with PA (250 ng/ml). (G) Primary BMDM kept at 37°C or 42°C for 1 h were then transfected with HT DNA (1 μg/ml) using Lipofectamine 2000 for 2 h. (H) iBMDM incubated at 37°C or at 41°C for 1 h followed by recovery at 37°C for 0-4 h, were simultaneously stimulated with LPS (100 ng/ml) and nigericin (5 μM) at 37°C for 1 h. Cell free supernatants were analysed by LDH release assay (A, C, H) or by ELISA for IL-18 (E) or TNF (F). Supernatants and cell lysates were analysed by western blotting for cleavage and expression of Caspase-1, GSDMD, and β-actin (B, D). Average value of three technical replicates from N=3 independent biological replicates (symbols), floating bars (min to max) line at mean (A, H). Average of three technical replicates from N=5-6 donors indicated by symbols linked by lines, floating bars (min to max) line at median (C,E,F). Representative blots from N=2-4 biological replicates (B,D,G). Data were analysed by Two-way RM ANOVA with Šídák’s multiple comparisons test p= <0.05* p< <0.01**; < 0.001***.< 0.0001****.

To confirm that NLRP3 inflammasome formation is inhibited by high temperature we investigated ASC speck formation by microscopy. As shown in Figure 2A-B HMDM pre-treated at 41°C formed significantly less ASC puncta/specks in response to NLRP3 stimulation (LPS + nigericin) than cells kept at 37°C. Heated BMDM also displayed dramatically reduced levels of ASC specks following NLRP3 activation by LPS and nigericin (Fig. 2C-D). As multiple PRRs can form inflammasomes with ASC, we examined the effects of high temperature on additional inflammasome sensors. NAIP/NLRC4 inflammasome activation by addition of the needle protein Lfn-MxiH and protective antigen (PA) caused ASC speck formation in HMDM that was not significantly decreased by incubation at 41°C (Fig. 2A-B). In BMDM, NLRC4 activation by the needle protein Lfn-BsaK was partially reduced in heated cells (Fig. 2C-D). We also examined pyroptosis and IL-1β release (Fig. 2E-F) following NAIP/NLRC4 activation in HMDM. As expected in these experiments pyroptosis and IL-1β secretion in response to NLRP3 activation were significantly reduced in heated cells. However, in response to activation NAIP/NLRC4 by LFn-BsaL, pyroptosis was not significantly reduced in cells incubated at 41°C (Fig. 2E). As priming is required to induce pro-IL-1β, we also analysed NAIP/NLRC4 activation in LPS primed HMDM. There was a small but significant decrease in pyroptosis and IL-1β and TNF secretion after exposure of cells to 41°C (Fig 2E-F, Supp. Fig. 2F). We also examined the effect of high temperature on activation of the double stranded DNA sensor AIM2. In BMDM, incubation at 42°C resulted in a small decrease in AIM2-dependent pyroptosis and GSDMD cleavage (Fig. 2G, Supp. Fig 2G). Together these data show that high temperatures can partially inhibit the NLRC4 and AIM2 inflammasomes, while NLRP3 is exquisitely sensitive to high temperature-mediated inhibition. As both NAIP/NLRC4 and AIM2 activation cause high levels of pyroptosis in these conditions (Fig. 2F, G), it is possible that K^+^ efflux caused by pyroptosis causes indirect NLRP3 activation which is affected by high temperature.

To address the mechanism of how high temperature regulates NLRP3 we tested whether we could recapitulate the effects of temperature in human embryonic kidney (HEK)293T cells that do not basally express inflammasome proteins. We used HEK293T lines stably expressing ASC-EGFP and HA-tagged NLRP3 or NLRP1 as previously described^33^. The addition of nigericin to NLRP3 expressing cells triggered ASC speck formation measured by flow cytometry (Fig. 3A). Incubating the cells at 41°C prior to the addition of nigericin caused a significant decrease in ASC specks (Fig. 3A). Val-boroPro (VbP, Talabostat) triggered ASC speck formation in NLRP1 expressing cells which was not inhibited by incubation at high temperature (Fig. 3B). These data demonstrate that high temperature specifically affects NLRP3 activation in an overexpression system, suggesting that NLRP3 itself is sensitive to high temperatures.

**Figure 3.**
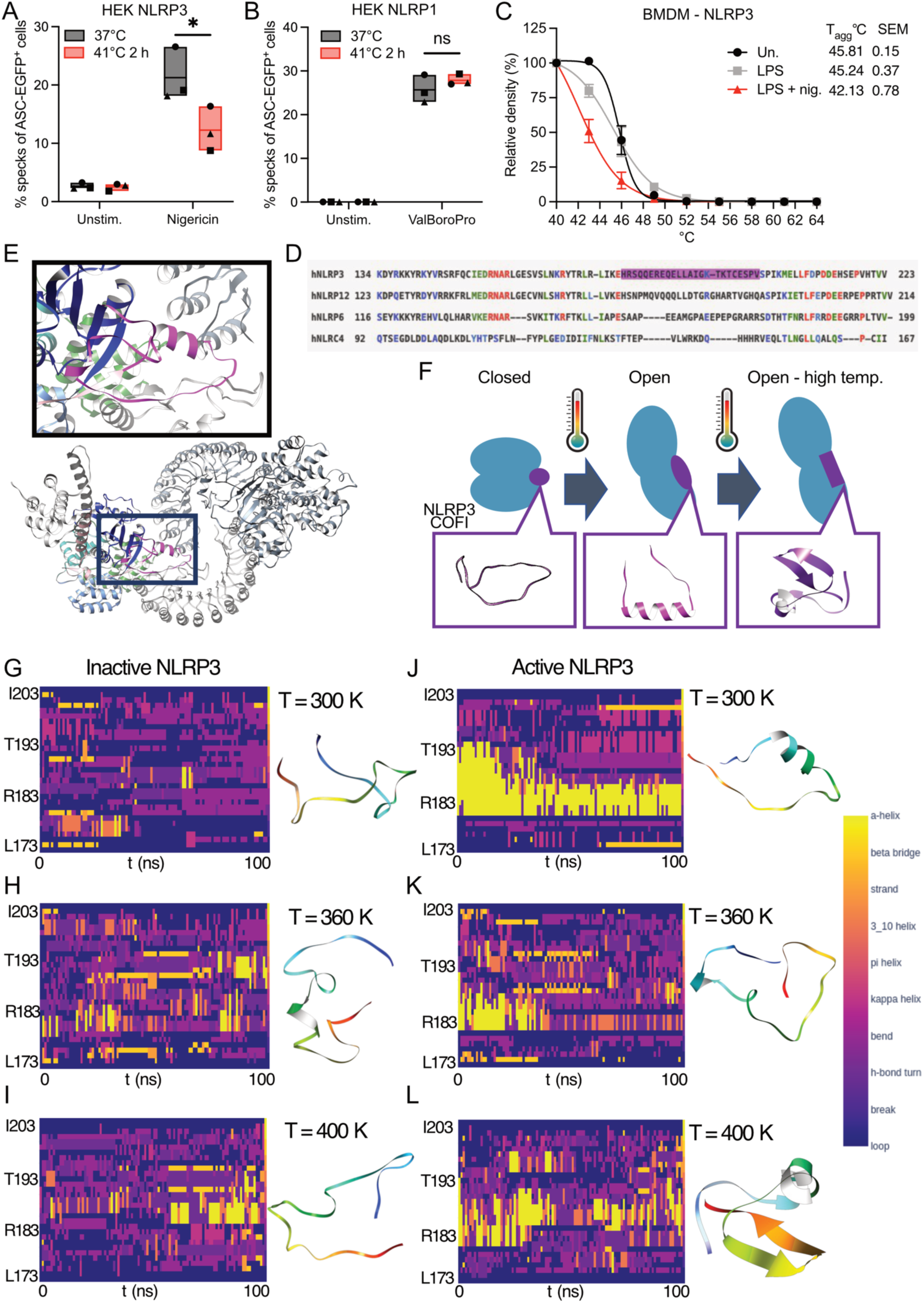
NLRP3 protein stability is highly sensitive to elevated temperature*s in cellulo* and *in silico*. (A-B) HEK293 cells stably expressing ASC-EGFP and NLRP3 or NLRP1 were incubated at 41°C for 3 h before stimulation with nigericin (5 μM, 1 h) or Talabostat (30 μM, 24 h) at 37°C. ASC specks were quantified by flow cytometry. (C) BMDM stimulated with LPS (100 ng/ml) for 3 h followed by nigericin (5 μM) in the presence of a caspase-1 inhibitor. CETSA was performed (40-64°C) and samples analysed by western blot for NLRP3. Relative density was calculated using ImageJ. Average value of from N=3 independent biological replicates (symbols), floating bars (min to max) line at mean (A, B). Average +/-S.E.M, of NLRP3 expression from N=4 independent biological replicates (C). Data were analysed by Two-way RM ANOVA with Šídák’s multiple comparisons test p= <0.05*. (D) Sequence alignment of FISNA subdomains of human NLRP3, NLRP6, NLRP12 and NLRC4. Conserved (red), highly similar (green), and weakly similar (blue) residues; NLRP3 COFI region (magenta). (E) Steric clash between the helical conformation of COFI and the LRR domain, and with the LRR domain of another interacting NLRP3 protein in the auto-inhibited “cage” oligomer (PDB code: 7PZC). NLRP3 domain/region colours: PYD (dark grey), FISNA (light pink), COFI (magenta), NBD (dark blue), HD1 (sky blue), WHD (marine green), HD2 (dark green) and LRR domain (light grey). (F) NLRP3 activation is coupled with NACHT domain (blue) adopting an open conformation and with loop-to-helix conformational change of the COFI (purple/magenta) facilitated by a mild increase in temperature. Further increase in temperature causes COFI to adopt an aggregation-prone conformation. (G-L) Changes in the per-residue secondary structure of the COFI induced by increased temperature for inactive (G-I) and active (J-L) NLRP3. Simulation time (X-axis) and residue number (Y-axis). The ribbon structure of the highest-populated cluster for each simulation is also shown, coloured by a rainbow gradient from N-(blue) to C-terminus (red).

**Supplemental Figure 3.**
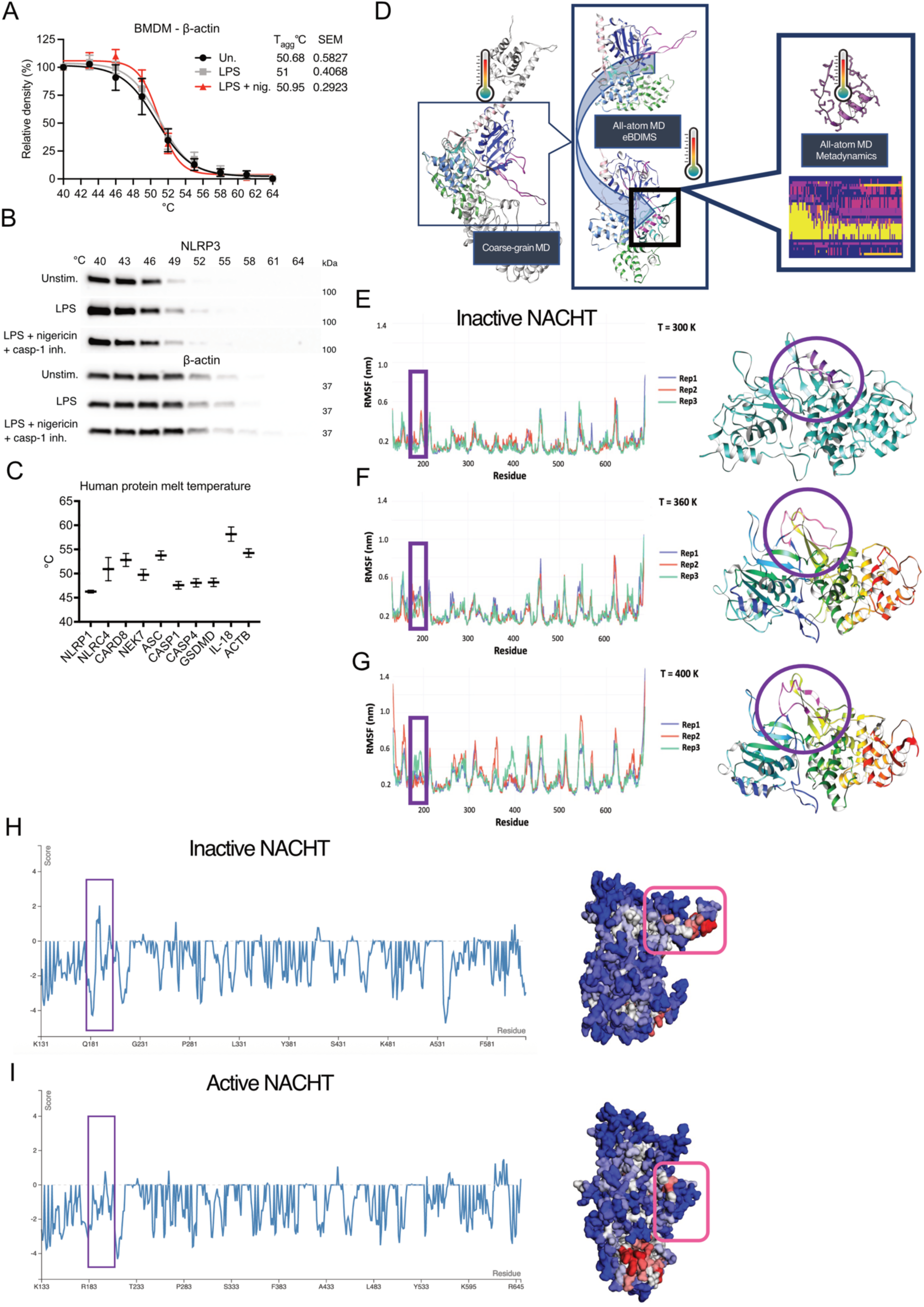
NLRP3 protein stability is highly sensitive to elevated temperature*s in cellulo* and *in silico*. (A-B) BMDM stimulated with LPS (100 ng/ml) for 3 h followed by nigericin (5 μM) in the presence of a caspase-1 inhibitor. CETSA was performed (40-64°C) and samples analysed by western blot for NLRP3 and β-actin. Relative density was calculated using ImageJ. Average +/-S.E.M, of β-actin expression from N=4 independent biological replicates (A), representative blots (B). (C) Melt temperatures of indicated proteins taken from the human cell line meltome dataset from Jarzab et al.^1^ Values are average +/-S.D. from N=4-35 measurements in human cell lines. (D) The multiscale workflow employed in this study (left to right): Fast coarse-grain (CG) molecular dynamics (MD) simulations of ΔPYD human NLRP3 were used to gain an insight into protein dynamics at microseconds time scale. These results were used to seed slower but more accurate all-atom MD simulations of the whole NACHT domain including the COFI region, and the data were used to “seed” Brownian Dynamics simulations (eBDIMS), to gain an insight intro conformational changes associated with NLRP3 activation. Finally, atomistic MD simulations of the COFI were used to assess the secondary structure content at a range of temperatures, and to carry out well-tempered metadynamics enhanced sampling simulations to gain more detailed understanding of COFI dynamics at high temperatures. (E-G) Conformational changes, quantified as root-mean-square fluctuations (RMSF) per-residue for closed conformation (inactive) NACHT domains at the standard temperature 300 K (E), high temperature 360 K (F), and very high temperature 400 K (G). Data for three independent simulation replicas are shown with representative 3D structures on the right panels. COFI region is highlighted in purple. (H-I) Aggrescan3D plots, showing the calculated propensity to aggregate (score above cutoff) and their corresponding 3D structures, showing predicted aggregation “hotspots”, low aggregation propensity shown in blue or high propensity shown in red. (H) NLRP3 NACHT domain in the inactive closed conformation, with the conformation of the COFI (purple box) being more extended and aggregation-prone. (I) NLRP3 NACHT in open active conformation, with the COFI (purple box) in the α-helical conformation.

To examine the effect of high temperatures on NLRP3 stability we next used a cellular thermal shift assay (CETSA). CETSAs determine the temperature at which 50% of a cellular protein will aggregate or ‘melt’ and can be used to examine small molecule binding, PTMs, and protein-protein interactions (PPIs)^34^. In our CETSA experiments we found that in unstimulated BMDM the temperature of aggregation (T_agg_) of NLRP3 was 45.81°C (Fig. 3C and Supp. Fig. 3B). In the same conditions the T_agg_ of β-actin was 50.7°C (Supp. Fig. 3A-B). A previous study using mass-spectrometry CETSA, also known as thermal proteome profiling, determined the melting temperatures (Tm) of >5000 human proteins in several cell lines^1^. NLRP3 was not observed in the dataset as it is not expressed in any of the cell lines that were tested. However, the melting temperatures of several inflammasome sensors and associated proteins including NLRP1 (46.26 °C), NLRC4 (50.91°C), ASC (53.74°C), caspase-1 (47.57°C), and NEK7 (49.74°C) were determined (Supp. Fig. 3C). NLRP3 thus appears to have a lower melting temperature than ASC, NEK7, and caspase-1 that are required to form the NLRP3 inflammasome signalling complex. We also examined if the activation status of NLRP3 affected its stability by stimulating BMDM with LPS and nigericin in the presence of a caspase-1 inhibitor to prevent pyroptosis (Fig. 3C, Supp. Fig. 3B). Priming did not significantly affect NLRP3 stability with NLRP3 from LPS stimulated cells having a T_agg_ of 45.24°C. However, activation shifted the T_agg_ of NLRP3 to 42.13°C suggesting that the active state of NLRP3 is considerably less stable than inactive or primed NLRP3. This is surprising as formation of NLRP3 oligomers and interaction with ASC were expected to stabilise NLRP3 through PPIs but suggests that upon activation NLRP3 conformational changes may destabilise NLRP3.

To complement our *in cellulo* analyses we next used a multiscale *in silico* approach to model the effects of high temperatures on NLRP3 intrinsic dynamics and structural stability (Supp. Fig. 3D). NLRP3 contains three domains: the N-terminal pyrin domain (PYD), the nucleotide-binding and oligomerisation domain known as the NACHT (domain present in NAIP, CIITA, HET-E and TP1), and the C-terminal leucine rich repeat (LRR) domain^11,35^. The central NACHT is made up of five subdomains, denoted as FISNA (fish-specific NACHT-associated), NBD (nucleotide binding domain), HD1 (helical domain 1), WHD (winged helix domain) and HD2, respectively. The FISNA was previously included within the NACHT, however it is now considered a separate subdomain^36^. The NLRP3 FISNA (residues 130-220 in human NLRP3) is unique and most NLRs do not contain such a region. The presence of the FISNA domain in NLRC4 and NAIP5 has been noted, however these proteins do not contain the C-terminal region (residues 176-202) present in NLRP3’s FISNA^37^.The N-terminal region of the FISNA, including a flexible linker connecting the PYD and FISNA, has been shown to be a key region for NLRP3 activation^37,38^, however the <u>C</u>-terminal <u>o</u>f <u>FI</u>SNA (COFI) region, also denoted as helix 2 in some works^39^, has been largely overlooked and is not conserved in closely related NLRs (Fig. 3D). Previous structural data identified a stable conformation of the FISNA in activated NLRP3 that is distinct from its conformation in auto-inhibited NLRP3^39^. This indicates there is a prominent role of the intrinsic dynamics of this region in the activation process. The activation of NLRP3 was associated with the ordering of the COFI, while the region (magenta) clashed with the LRR domain (grey) in the closed/auto-inhibited conformation (Fig. 3E). However, details of conformational transitions coupled with the ordering have not been explored.

Conformational transitions in large, multidomain proteins such as NLRP3 are cooperative processes with many short-lived intermediates. These intermediates are often inaccessible to structural biology techniques but can be captured by molecular simulations. To get an insight into the intrinsic dynamics of the NLRP3 FISNA and its conformational changes during the transition from the closed to open conformation, we combined molecular dynamics simulations at standard (300 K), and heated (360-400 K) conditions, and used these to “seed” coarse-grain Brownian dynamics simulations to detect conformational transitions between “closed” (auto-inhibited) and “open” (activated) conformations of the NACHT domain. This was supplemented by all-atom equilibrium MD simulations at different temperatures and enhanced sampling calculations using well-tempered metadynamics (Supp. Fig. 3D). We observed that the COFI is likely to be highly sensitive to temperature changes, and data analysis combined with the Aggrescan3D tool indicated it is aggregation-prone (Fig. 3F-L, Supp. Fig. 3E-I).

As the COFI region is absent from the experimental structure of closed/auto-inhibited conformations of NLRP3, it had to be modelled. AlphaFold models the COFI in a predominantly helical conformation, as observed in the active NLRP3 disc, but this is not compatible with the auto-inhibited (closed) conformation of NLRP3^39^. Therefore, we modelled the isolated COFI forcing an extended conformation compatible with the steric requirements of the auto-inhibited NLRP3 dodecamer (PDB code: 7PZC) and used the model as a starting point of our investigations of COFI’s intrinsic dynamics and temperature dependence. We found that increasing the temperature promoted an open conformation of the NLRP3 NACHT domain; a necessary step preceding the activation of NLRP3. This conformational change is coupled with a loop-to-helix transition of the COFI (Fig. 3F, Supp Fig. 3E-F, Supp. Movie 1). However, in the active (open) conformation at increased temperature, the helix unfolded and transitioned to an aggregation-prone β-sheet-like conformation (Fig. 3F, Supp. Movie 2A). This conformational change may therefore contribute to diminished NLRP3 activity at high temperature. At the highest temperature (400 K) we observed signs of early-stage thermal unfolding, but up to 360 K the NACHT domain remained folded, with the major conformational changes occurring in the COFI region (Supp. Fig. 3E-G). We next carried out MD simulations of the isolated COFI region at temperatures spanning from 300 K to 400 K and used the highest-populated clusters of those simulations to seed well-tempered metadynamics simulations (Fig. 3G-L). Unexpectedly, the inactive closed conformation of the NACHT domain was stabilized through additional residue contacts between the COFI and the remaining protein core at moderately increased temperature (330 – 360 K). At these temperatures, the COFI’s secondary structure content shifted towards more helical (Fig. 3G, H). At the highest temperature (400 K), COFI adopted aggregation-prone coil conformation, which is consistent with signs of thermal unfolding (Fig. 3I, L).

To understand the functional importance of the COFI, we evaluated a COFI null mutant, wherein residues 177-195 were removed, and a variant wherein the COFI region was replaced by the homologous region of the same length derived from closely related NLRP12 (Fig. 4A-D, Supp. Fig. 4A-C). Modelling NLRP3Δ177-195 resulted in the protein being less prone to aggregation at increased temperatures, yet it retained all the key features of wild-type NLRP3 NACHT in terms of ability to adopt two distinct conformations and retained binding sites for nucleotides and MCC950. However, unexpectedly the deletion of COFI favoured the closed conformation of the NACHT at all temperatures, due to additional interactions between the solvent-exposed portion of the LRR (residues 708-722) and NACHT in the absence of COFI (Supp. Fig 4B). Compared to the MD simulation of wild-type NLRP3, the NLRP3Δ177-195 monomer was more stable overall. The deletion of the aggregation-prone COFI caused the COFI-null NLRP3 to show less per-residue root-mean-square fluctuations (RMSF) and lower backbone root mean square deviation (RMSD), indicating lower protein stability (Supp. Fig. 4E-H). On the other hand, the NLRP12 COFI swap mutant showed intrinsic dynamics features that are distinct from wild-type NLRP3 and NLRP3Δ177-195. The NLRP12 COFI mutant promoted the closed conformation of the NACHT domain via increased stabilisation of helix-helix interactions between the “swapped” region and residues from the LRR domain, effectively “glueing” the NACHT and LRR (Supp. Fig. 4C). These data support the hypothesis that the COFI is strongly linked to conformational changes in the NLRP3 monomer that are required for NLRP3 activation.

**Figure 4.**
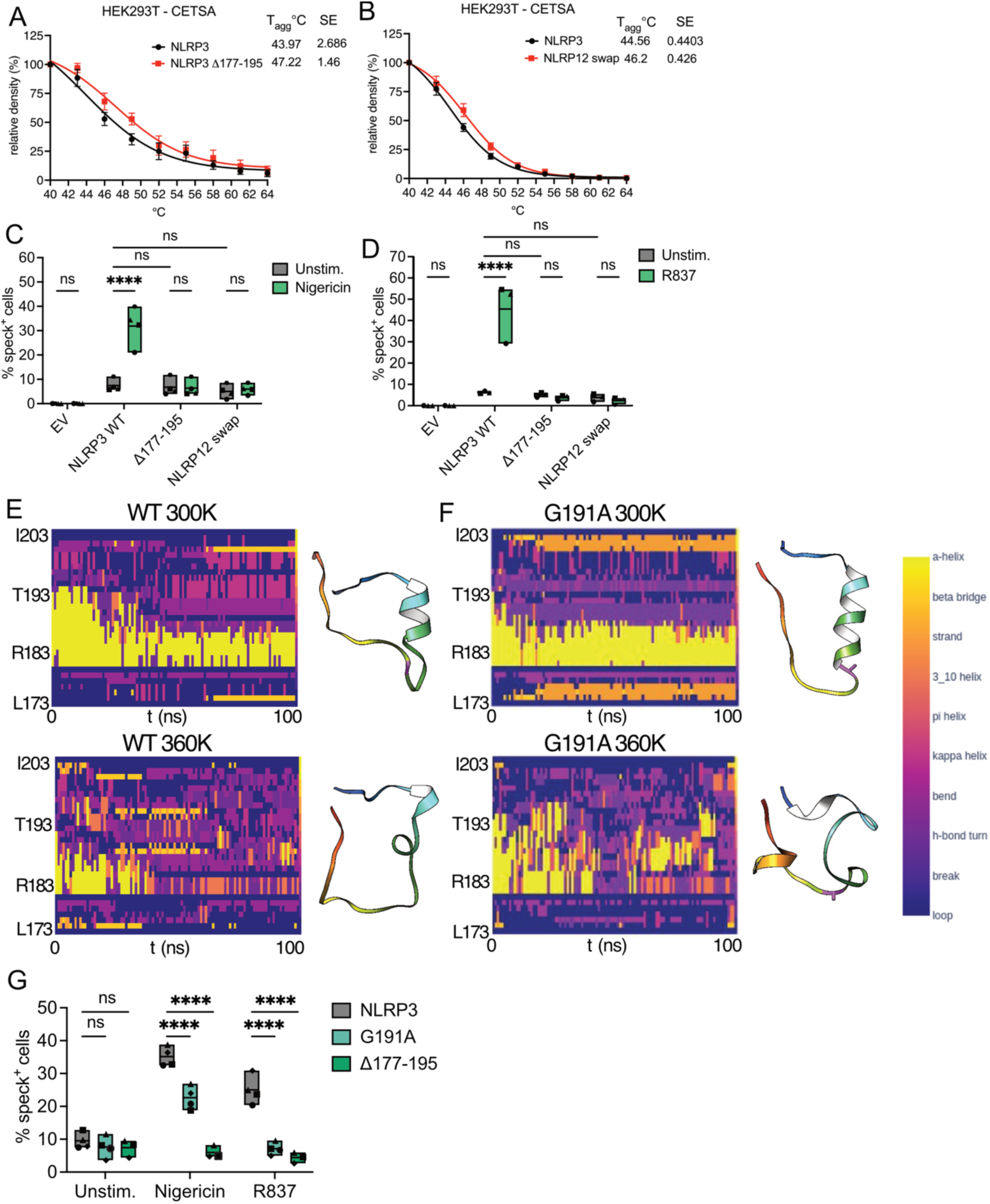
The NLRP3 COFI senses changes in temperature and is required for NLRP3 activation. (A-B) HEK293T cells transfected with plasmids encoding wild-type human NLRP3, NLRP3β177-195, or NLRP3-NLRP12 COFI swap mutants. CETSA was performed (40-64°C) and samples analysed by western blot for NLRP3. Relative density was calculated using ImageJ. Average +/-S.E.M of NLRP3 expression from N=6 (A) and N=3 (B) independent biological replicates. (C,D,G) HEK293T cells stably expressing hASC-GFP transfected with plasmids encoding cherry tagged-wild-type human NLRP3, NLRP3β177-195, NLRP3-NLRP12 COFI swap, NLRP3 G191A, or empty vector control. Cells were unstimulated or were treated with nigericin (5 μM, 1 h) or R837 (100 μM, 4 h). ASC specks quantified by flow cytometry. (C, D, G) Average of N=3-4 independent biological replicates (symbols), floating bars (min to max) line at mean. Data were analysed by Two-way RM ANOVA with Šídák’s multiple comparisons test p=< 0.0001****. (E) Changes in the per-residue secondary structure of the COFI region (open conformation of NLRP3 NACHT domain) induced by increased temperature for G191A (upper panels) and wild-type NLRP3 (lower panels). Simulation time (X-axis) and residue number (Y-axis). The ribbon structure of the highest-populated cluster for each simulation is also shown, coloured by a rainbow gradient from N-(blue) to C-terminus (red). Residue 191 is coloured purple.

**Supplemental Figure 4.**
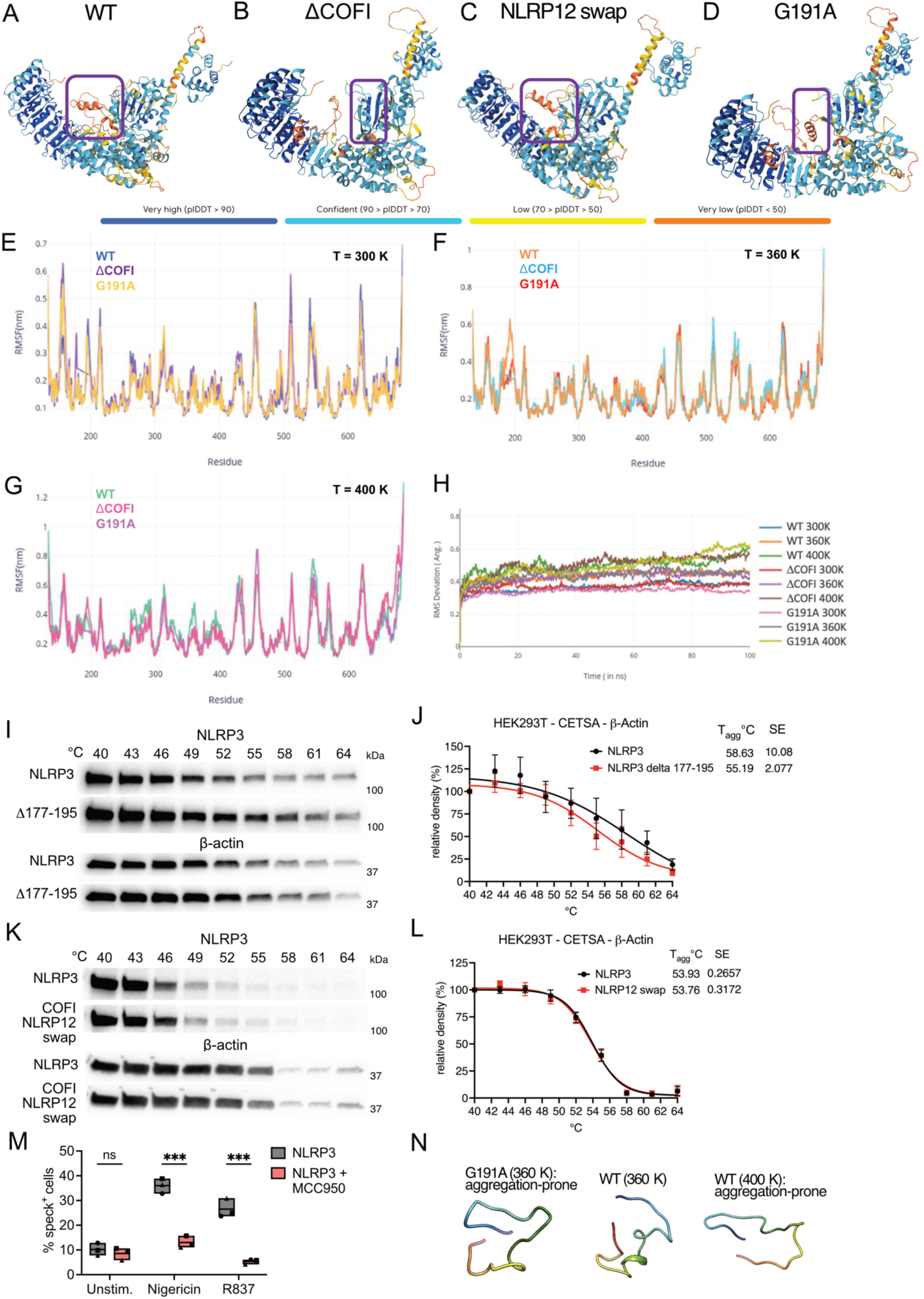
T**h**e **NLRP3 COFI senses changes in temperature and is required for NLRP3 activation.** (A-D) Top-ranking AlphaFold3 models of human wild-type (WT) NLRP3 (A), COFI deletion β177-195 (B), NLRP3-NLRP12 COFI swap mutant (C) and G191A mutant (D). Models are coloured by confidence score. Although the COFI region is consistently modelled in the α-helical conformation in WT and G191A, the orientation of the helix relatively to the NACHT and LRR domains are different for both proteins. In the βCOFI mutant, the modelled arrangement of NACHT domain is similar to WT, and the region connecting FISNA and NACHT is modelled with high-confidence. In NLRP3-NLRP12 COFI swap mutant the COFI is consistently modelled in the α-helical conformation but the relative orientation of the α-helix differs, and the swap mutant is stabilised by an additional β-sheet segment. (E-G) Conformational changes, quantified as root-mean-square fluctuations (RMSF) per-residue for open conformation NACHT domains of wild-type NLRP3 (WT), COFI deletion mutant (ΔCOFI), and G191A mutant at standard temperature 300 K (E), high temperature 360 K (F), and very high temperature 400 K (G). (D) Root-mean-square deviations (RMSD) for each temperature, three independent simulation replicas. (I-L) HEK293T cells transfected with plasmids encoding wild-type human NLRP3, NLRP3β177-195, or NLRP3-NLRP12 COFI swap mutants. CETSA was performed (40-64°C) and samples analysed by western blot for NLRP3 and β-actin. Relative density was calculated using ImageJ. Average +/-S.E.M of β-actin expression from N=6 (F) and N=3 (H) independent biological replicates, (I, K) representative blots. (M) HEK293T cells stably expressing hASC-GFP transfected with a cherry tagged-wild-type human NLRP3 plasmid. Cells were pre-treated with MCC90 (100 nM, 1 h) and left unstimulated or were treated with nigericin (5 μM, 1 h) or R837 (100 μM, 4 h). ASC specks were quantified by flow cytometry. Average value from N=3 independent biological replicates (symbols), floating bars (min to max) line at mean. Data were analysed by Two-way RM ANOVA with Šídák’s multiple comparisons test p=< 0.001***. (N) Structures of the highest-populated clusters for each temperature simulation (closed inactive NLRP3 NACHT), coloured by rainbow gradient from N-(blue) to C-terminus (red).

To examine the function of the COFI in cellular assays, we first examined whether deletion of the COFI or swapping in the NLRP12 sequence affected NLRP3 stability in CETSAs. We expressed wild-type human NLRP3, NLRP3Δ177-195 or the NLRP12 COFI swap in HEK293T cells. CETSA showed that the T_agg_ of overexpressed NLRP3 was ∼44°C, while NLRP3Δ177-195 (47.22°C) and NLRP12 swap (46.2°C) were more stable (Fig. 4A-B, Supp. Fig. 4I, K). In the same assays the T_agg_ of β-actin was between 53.76-58.63°C (Supp Fig. 4I-L), in line with the melt temperature of human β-actin determined in previous TPP datasets (54.26°C -Supp. Fig. 3C). This suggests that the COFI may be involved in destabilising NLRP3 at high temperature in agreement with our molecular simulations. We next examined the involvement of the COFI in NLRP3 activation using an ASC speck formation assay. HEK293T cells stably expressing ASC-GFP were transfected with plasmids to express wild type human NLRP3 tagged with mCherry. ASC speck formation was monitored by dual colour time-of-flight inflammasome evaluation (TOFIE) flow cytometry technique as previously described^40,41^. As expected, expression of wild type NLRP3 triggered some ASC speck formation (∼10%) in unstimulated cells (Fig. 4C-D). However, the addition of nigericin or R837 significantly enhanced ASC speck formation (Fig. 4C-D). Importantly, MCC950 treatment blocked nigericin and R837 induced

ASC speck formation confirming that NLRP3 activation can be examined using this approach. (Supp Fig. 4M). When we expressed the NLRP3Δ177-195 or NLRP12 swap mutants some ASC speck formation was induced in unstimulated cells similar to wild-type NLRP3. However, treatment with nigericin or R837 failed to increase ASC specks in the NLRP3Δ177-195 or NLRP12 swap mutants (Fig. 4C-D). This suggests that the COFI is necessary for stimulus-induced NLRP3 activation, in agreement with our molecular modelling and simulations.

Through our NLRP3 simulations and Aggrescan3D analysis (Supp. Fig. 3H-I), we identified the G191 residue within the COFI as a “hotspot” for helix-to-coil conformational transition and for aggregation. This is consistent with reported effects of glycine residues on α-helices, where glycines are known as α-helix destabilisers when positioned in the middle of the helix but favoured at the flanking regions of the helices due to their small size and high flexibility^42^. We investigated an NLRP3 G191A mutant, as alanine is a known helix-stabiliser at standard temperatures. For G191A, the RMSF analysis showed a similar fluctuation pattern of the residues at the whole temperature range (300-400 K) (Supp. Fig. 4E-H). It is noted that the COFI had an active fluctuation at all the simulated temperatures. As the temperature increased, the RMSF values in the COFI increased from 0.5 nm to 1.0 nm. Although a mild increase was observed in the region of residues 180-199, where conformational changes are observed in wild-type NLRP3, we hypothesised that the presence of alanine restricted the conformational space and caused a hinderance in the amount of fluctuation, therefore preventing a conformational change (Fig. 4E-F, Supp. Movie 2B). Simulations of G191A in the closed conformation of the NACHT domain showed unexpected stabilisation of the closed conformation at higher temperatures, due to enforcing different conformations of the COFI distinct from wild type NLRP3 and increased contacts with NACHT and LRR domains (Fig. 4E-F, Supp. Fig. 4D, N). In an ASC speck formation assay, the G191A mutant had significantly reduced response to both nigericin and R837, similar to the NLRP3Δ177-195 mutant (Fig. 4G). Together both our cell-based experiments and *in silico* observations identify the NLRP3 COFI region and G191 residue as critical for NLRP3 activation and for regulating NLRP3 stability at high temperature.

As our data demonstrated that FRT could significantly inhibit NLRP3-dependent inflammation we asked if FRT also affects inflammatory responses *in vivo*. We used a model developed by Munoz et al.^43^ to increase the core body temperature of mice. Mice placed in a heated chamber 38°C for 18 hours had a 1.72°C increase in core temperature measured by rectal thermometer compared to standard housing temperature (22°C) (Fig. 5A-B). Following 2 h of recovery at 22°C mice were challenged with LPS by intraperitoneal (IP) injection for 2 h. In Figure 5C-D, analysis of serum cytokine levels showed that there was no significant change in levels of IL-1β and IL-18 in the heat exposed mice, although there was a trend to lower levels. However, there were significant reductions in IL-6, TNF, MCP-1 (CCL2) and RANTES (CCL5), demonstrating that high body temperature reduced inflammation in response to LPS *in vivo* (Fig 5E-H). Importantly, not all cytokines were altered in heat exposed mice, with levels of GM-CSF, IL-12 p70 and MIP-1α/β all unchanged between the groups (Supp. Fig 5A-D). The IP LPS challenge model has previously been shown to be NLRP3-dependent^27,44^. While we did not see a significant effect on IL-1β and IL-18, we did see reductions in cytokines that are induced by IL-1 (IL-6, TNF, MCP-1 and RANTES)^45–47^. It is thus possible that high temperature reduced LPS-induced IL-1β at an earlier timepoint. Of particular interest, the anti-inflammatory cytokine IL-10 was significantly increased in heat-exposed mice (Fig 5I). How IL-10 is regulated by temperature is currently unknown, but we note that IL-1β and IL-18 can negatively regulate IL-10 production in T cells^48–50^. These *in vivo* data demonstrate that high temperatures reduce inflammation in response to LPS and this may be mediated in part via the attenuation of NLRP3 signalling.

**Figure 5.**
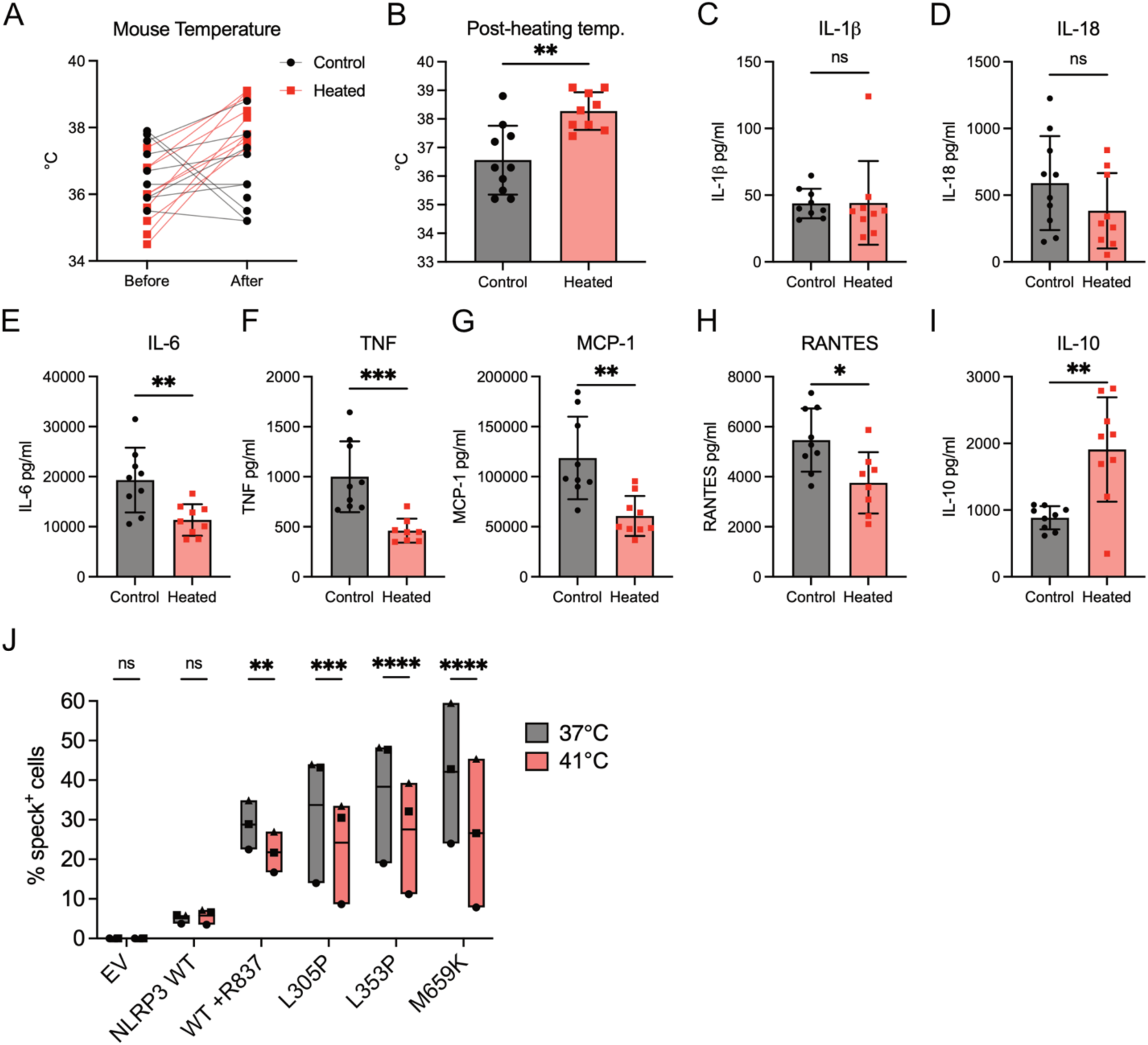
High temperature reduces LPS-induced inflammation *in vivo* and hyperactivation of FCAS-associated NLRP3. (A-B) Core body temperature measurements from mice housed at 38°C for 18 h (heated) or at 22°C (control). (C-I) Serum cytokine measurements from heat exposed and control mice 2 h after intraperitoneal LPS challenge. N=9 mice per group. Groups are compared using an unpaired two tailed t-test. (J) HEK293T cells stably expressing hASC-GFP transfected with plasmids encoding cherry tagged-wild-type human NLRP3 or FCAS-associated mutations. Cells were incubated at 37°C or at 41 °C for 3 h. WT NLRP3 was left unstimulated or was treated with R837 (100 μM, 4 h). FCAS mutants were unstimulated. ASC specks were quantified by flow cytometry. Average value from N=3 independent biological replicates (symbols), floating bars (min to max) line at mean. Data were analysed by Two-way RM ANOVA with Šídák’s multiple comparisons test. p= <0.05*, <0.01**; <0.001***, <0.0001****.

**Supplemental Figure 5.**
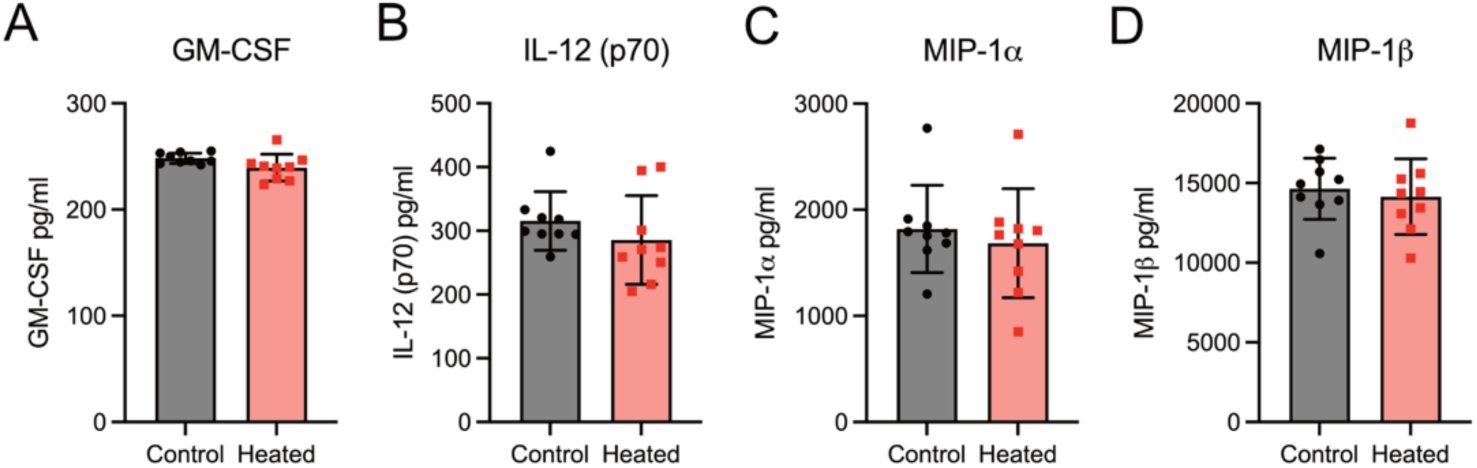
Effects of high temperature on LPS-induced inflammation *in vivo*. (A-D) Serum cytokine measurements from heat exposed and control mice 2 h after intraperitoneal LPS challenge. N=9 mice per group. Groups are compared using an unpaired two tailed t-test.

To understand whether our observations could have therapeutic implications, we examined how high temperature affects NLRP3 FCAS mutations (L305P, L353P and M659K) using the dual colour TOFIE approach. As expected, activation of NLRP3 with R837 increased ASC speck formation that was attenuated by incubation at high temperature (Fig. 5J). Expression of hyperactive FCAS NLRP3 resulted in high levels of ASC speck formation that were even greater than activated wild-type NLRP3. Strikingly, the number of ASC specks induced by the FCAS mutants was significantly reduced upon incubation at 41°C (Fig. 5J). These data show that disease-associated NLRP3 is also inhibited by high temperatures.

## Discussion

The return to homeostasis is the optimal outcome of the inflammatory response following infection or injury. However, the molecular details that underpin the resolution of inflammasome-driven inflammation are underappreciated. Herein we have discovered a novel mechanism for self-limiting regulation of NLRP3 inflammasome signalling that is mediated by one of the key outcomes of this pathway, elevated temperature. We have demonstrated that short term incubation for one hour at high FRT specifically blocks NLRP3 signalling but has limited impact on other inflammasomes and TLR4 signalling. Our findings are relevant to the systemic increase in temperature that occurs during fever, but also to localised inflammation where significant increases in temperature occur at the site of injury (calor)^51^. For example, a study from over a century ago demonstrated that following the fracture of a femur of a guinea pig the inflamed tissue was >3°C warmer than uninjured tissue^52^. More recent thermography studies have examined the potential of using temperature measurements of joints and tissues from rheumatoid arthritis patients to predict disease progression, while thermography may also aid in the assessment of wound infection^51,53–55^. Interestingly, it has been demonstrated *in vivo* that NLRP3 signalling is essential for increased local tissue temperatures, as *Nlrp3*^-/-^ mice have significantly lower ankle temperatures following intra-articular injection of monosodium urate (MSU) crystals^56^. The negative regulation of NLRP3 by high temperatures may thus help resolve localised NLRP3-driven inflammation at sites of infection and injury by preventing extended inflammasome activation and signalling.

While we have tested physiologically relevant temperatures that can occur at high FRT, our cell models do not recapitulate the sustained but generally milder elevated temperatures that occur during fever. Therefore, we attempted to model the effects of fever *in vivo* using a whole-body warming approach. While our model significantly increased mouse body temperatures and reduced LPS-induced inflammatory cytokines, it did not significantly reduce inflammasome-dependent IL-1β and IL-18 levels at the time point examined. The response to LPS *in vivo* is complex as LPS activates both TLR4 and the intracellular non-canonical inflammasome, that in turn indirectly activates NLRP3 via potassium efflux^57^. We observed relatively low levels of IL-1β release in this experiment relative to previous similar studies^43^, which may explain the apparent lack of effect of heating. A more specific NLRP3-dependent response could be examined in future using an LPS plus ATP-induced peritonitis model^58,59^. Nevertheless, whole-body warming clearly reduced serum levels of several IL-1-inducible cytokines and chemokines after LPS treatment *in vivo.* This is notable as fever and localised tissue temperature increases are generally associated with promoting the immune response, for example through enhancing leukocyte trafficking^3,51^. Recent studies have also demonstrated that FRT can specifically modulate mouse T cell metabolism leading to increased T cell activation, proliferation and effector functions^7,60^. This suggests that fever generally enhances the adaptive immune response during infection. In contrast, our data suggest that fever negatively regulates elements of the innate immune response to prevent excessive inflammation or a prolonged and extremely metabolically costly febrile response. Future work examining the impacts of temperature on innate immune cell metabolism may reveal differences in the metabolic adaptations to temperature stress in innate cell types.

Our findings have potential implications for understanding the anti-inflammatory effects of mild hyperthermia. Elevated body temperatures, such as those achieved during fever, heat therapy, or even sauna use, have been historically linked to improved immune regulation and reduced inflammation^51^. Interestingly, clinical observations have shown that patients with FCAS often report symptom relief when residing in or traveling to warmer climates^61,62^. These observations support our finding that FCAS mutant NLRP3 is significantly inhibited by high temperature. More broadly, this suggests that NLRP3-mediated diseases may benefit from mild hyperthermia. Indeed, mild hyperthermia has demonstrated neuroprotective and anti-inflammatory effects in models of Alzheimer’s disease^63,64^, where NLRP3 causes damaging inflammation^65^, supporting its therapeutic potential.

The concept of molecular thermosensors has been pioneered in the study of plants where DNA, RNA and protein thermosensors have been described that control responses cold and heat stress^66^. In animals, membrane associated transient receptor potential (TRP) channels that sense cold and heat are well characterised. However, they are largely expressed in sensory neurons and only TRPM2 has been linked to macrophage function^67^. Our observation that the COFI of human and mouse cytosolic NLRP3 regulates its activation and is sensitive to temperature is thus novel. There are several criteria that have been proposed to define a thermosensor; these are: that temperature directly alters properties like structure or activity, that these properties are important for signal transduction, and that these properties are important for temperature-dependent physiological read-outs^66^. Our identification of the COFI peptide as a key region mediating temperature sensitivity in NLRP3, and that temperature regulates NLRP3 inflammasome formation and signalling, demonstrates that NLRP3 is a bone fide thermosensor.

## Resource availability

Requests for further information and resources should be directed to and will be fulfilled by the lead contact, Rebecca Coll. All plasmids generated in this study will be made available on request, but we may require a payment and/or a completed materials transfer agreement if there is potential for commercial application. All data reported in this paper will be shared by the lead contact upon request. This paper does not report original code.

## Acknowledgements

The work in the Coll (RCC) lab was supported by the Biotechnology and Biological Sciences Research Council (BB/V016741/1), Academy of Medical Sciences (SBR005\1104), the Medical Research Council (MR/Y014065/1), and start-up funding from WWIEM/QUB. The work in the Bronowska (AKB) lab was supported by the BBSRC (BB/T008695/1; scholarship for AS) and EPSRC (EP/S022791/1). Work by MJR and MAM was supported by National Health & Medical Research Council grants 2017/GNT1139644 and 2024/GNT2036891. We thank Prof Florian Schmidt (University of Bonn) for sharing the HEK293 cell lines stably expressing hASC-EGFP and hNLRP3 or hNLRP1. We thank Prof Kate Schroder (University of Queensland, Australia) for sharing the pEF6 human NLRP3-mCherry construct and HEK293T cells stably expressing hASC-GFP. We also thank Gabor Horvath and the Microscopy Core Facility of the Medical Faculty at the University of Bonn for providing help, services, and devices funded by the Deutsche Forschungsgemeinschaft (DFG, German Research Foundation, project number 388158066).

## Author contributions

Conceptualization: RCC, WW, AKB. Funding acquisition: RCC, AKB, MJR, BSF. Investigation: WW, RCC, DB, JZ, MAM, SX, AS, CMM, RK, MC. Methodology: AKB, SX. Project administration: RCC, AKB, MJR, BSF. Supervision: RCC. Visualization: JZ, SX, AS, AKB. Writing – original draft: RCC, WW, AKB. Writing – review & editing: all authors

## Supplemental material

**Supplemental Movie 1. Intrinsic dynamics of NACHT domain backbone, revealed during 100 ns of equilibrium MD) simulation of the NACHT domain in closed (inactive) conformation**.

Sub-domains are coloured as follows: FISNA – sky blue, COFI – brick red, NBD – yellow, HD1 – green, WHD – pink, HD2 – rose. (A) Shows the trajectory at room temperature (300 K). (B) Shows the trajectory at high temperature (360 K), at which the COFI region shows a tendency to shift to a transient α-helix. The magnified region shows the conformational transition of the COFI, observed at the higher temperature.

**Supplemental Movie 2. Intrinsic dynamics of NACHT domain backbone, revealed during 100 ns of equilibrium molecular dynamics (MD) simulation of the NACHT domain in open (active) conformation.**

Sub-domains are coloured as follows: FISNA – sky blue, COFI – brick red, NBD – yellow, HD1 – green, WHD – pink, HD2 – rose. The COFI is a helix in the open conformation. (A) Shows the COFI of wild-type human NLRP3 undergoing unfolding at higher temperature and adopting an aggregation-prone conformation. (B) Shows dynamics of the NLRP3 G191A mutant, at higher temperature, with the COFI remaining more stable and retaining a helical conformation. The side chain of the mutated residue (A191) is shown.

## Material and Methods

### Isolation and culture of primary human monocyte-derived macrophages (HMDM)

Buffy coats were obtained from the Northern Ireland Blood Transfusion Service (NIBTS reference number 2019/05), ethical approval for the use of buffy coats was from the Queen’s University Faculty Research Ethics Committee (reference MHLS 19_17). Human PBMCs were obtained from buffy coats by density gradient centrifugation in Ficoll–Paque PLUS. PBMCs were incubated at 4°C with CD14 magnetic microbeads and isolated by positive magnetic selection according to manufacturer’s instructions (Miltenyi Biotec). Macrophages were generated through differentiation of CD14^+^ monocytes with recombinant human M-CSF 50 ng/mL (for 7 days) or 100 ng/mL (for 3 days) in HMDM medium: RPMI 1640 medium containing 10% FBS, 1% penicillin/streptomycin (P/S), 1 × GlutaMAX (Thermo Fisher, 35050038), and 1 mM sodium pyruvate (Life Technologies, 11360070) at 37°C in a humidified 5% CO_2_ incubator. HMDM were harvested by scraping in HMDM medium, were rinsed with cold PBS, and then seeded onto 96-well plates at 0.7 × 10^5^ in 100 μL per well in HMDM medium.

### Culture of primary mouse bone marrow-derived macrophages (BMDM)

Primary bone marrow-derived macrophages from C57BL/6J mice housed at Queen’s University Belfast were generated as previously described^68^. Progenitors were differentiated for 7 days in BMDM medium: RPMI 1640 containing 10% FBS, 1% P/S and 1× GlutaMAX, with 50 ng/mL human M-CSF at 37°C in a humidified 5% CO_2_ incubator. Differentiated BMDMs were harvested with cold PBS and were plated onto 96-, 24-, and 6-well plates at 1 × 10^5^ in 100 μL, 2 × 10^5^ in 500 μL, and 1 × 10^6^ in 1000 μL cells per well respectively in BMDM medium.

### Culture of mouse immortalized bone marrow macrophages (iBMDM)

C57BL/6 iBMDMs (BEI Resources) were grown in iBMDM medium: DMEM containing 10% FBS, 1% P/S and 1 mM sodium pyruvate at 37°C in a humidified 5% CO_2_ incubator. Cells were passaged using trypsinisation and were routinely tested for Mycoplasma. iBMDMs were seeded onto 96-well plate at 0.5 × 10^5^ in 100 μL per well in iBMDM medium.

### Culture of human induced pluripotent stem cell (iPSC)-derived macrophages (iMacs)

KOLF2-C1 iPSCs were obtained from the Wellcome Sanger Institute (Hinxton, UK). iPSCs were cultured in StemFlex medium (ThermoFisher, A3349401) on vitronectin-coated (ThermoFisher, A14700) plates at 37°C in a humidified 5% CO_2_ incubator. 10 μM Rho-kinase (Rock) inhibitor Y-27632 Dihydrochloride (Merck, Y0503) was added for the first 24 hours following thawing and passaging. Embryoid bodies (EBs) were formed as previously described^69^. In brief, iPSCs were harvested using 5 mM PBS-EDTA (ThermoFisher, 15575020) and resuspended in StemFlex medium supplemented with 50 ng/mL BMP-4 (R&D Systems, 314-BP010) and 10 μM Rock inhibitor. iPSCs were seeded onto low-adherent, U-bottom 96 well plates at 1 × 10^5^ cells in 100 μL per well. The 96 well plates were centrifuged for 3 min at 300 × g, and incubated for 5 days to form EBs. On day 2 and day 4, 50 µL medium was removed and replaced with fresh medium. On day 5, the EBs were transferred from 96 well plates to gelatine-coated (Sigma-Aldrich, G1890) tissue culture flasks in myeloid precursor base medium: X-VIVO 15 Serum Free Medium (Lonza, BE02-060F), 1 × GlutaMAX and 1% P/S. Media was changed every 4–5 days. Approximately 3–4 weeks after transferring the EBs to the gelatine-coated plates, they began to produce myeloid precursor cells. To harvest the myeloid precursor cells, the medium was removed from the EB flasks and filtered through a 70 μm cell strainer. The filtered medium was centrifuged for 3 min at 300 × g and the pellet resuspended in iMac medium: RMPI 1640, 10 % FBS, 1 × GlutaMAX and 1% P/S supplemented with 100 ng/mL M-CSF. Myeloid precursor cells were plated at 5 × 10^6^ cells per plate and differentiated for 7 days. Differentiated iMacs were harvested using Lidocaine monohydrate (Sigma-Aldrich, L5647) and were plated onto 96 well plates at 0.5 × 10^5^ in 100 μL cells per well respectively in iMac medium.

### Inflammasome activation in macrophages

To activate the NLRP3 inflammasome in BMDMs and iBMDMs, LPS (100 ng/mL) was used to prime the cells for 3 h, followed by the addition of nigericin (5 μM), R837 (100 μM), or ATP (2 mM) for 1 h.

To activate the AIM2 inflammasome in BMDMs, LPS (100 ng/mL) was used to prime the cells for 4 h, and cells were transfected with 1 μg/mL herring testes DNA (D6898, Sigma-Aldrich) using Lipofectamine 2000 for 2 h. To activate the NLRP3 and NLRC4 inflammasome in HMDMs and iMacs, LPS (100 ng/mL) was used to prime the cells for 3-4 h, followed by the addition of Nigericin (5 μM) or Bacillus anthracis protective antigen (PA, 250 ng/mL) + Lfn-needle (1 ng/mL) for 2 h, respectively.

### Lactate Dehydrogenase (LDH) assay

Cells were plated in 96 well plates in culture medium overnight. Medium was then replaced with 100 μL Opti-MEM per well. Inflammasome stimuli were added for times indicated and at the end of the experiment 2 μL 10% Triton X-100 solution was added to the 100% lysis control wells. Plates were centrifuged for 3 min at 500 × g. Supernatants were removed and 25 μL supernatant was added to 25 μL LDH assay buffer (Cytotoxicity Detection Kit Roche, Sigma Aldrich, 11644793001 or LDH-Glo Cytotoxicity Assay, Promega, J2381) in a transparent 96 well plate. The plate was incubated in the dark for 30 min (mouse cells) or 1 hour (human cells) and absorbance was measured at 490 nm using FLUOStar Omega plate reader (BMG Labtech).

### Inflammasome assay cytokine measurement

Cytokine levels in cell-free supernatants were analysed by ELISA. Mouse IL-1β (Invitrogen, 88-7013-77), mouse TNF-α (Invitrogen, 88-7324-77; R & D Systems, DY410), human IL-1β (Invitrogen, 88-7261-77; R & D Systems, DY201), human TNF-α (Invitrogen, 88-7346-77; R & D Systems, DY210), human IL-6 (Invitrogen, 88-7066-88), and human IL-18 matched antibody pair (Invitrogen, BMS267-2MST) were used according to the manufacturer’s instructions.

### Western blotting

Cells were lysed in buffer (66 mM Tris HCl pH 7.4% SDS) with 2.5 Units BaseMuncher (Expedeon, BM0025). For some experiments protein concentration was determined using the PierceTM Rapid Gold BCA Protein Assay Kit (ThermoFisher, A53226) according to manufacturer’s instructions. 1 × NuPAGE LDS sample buffer (Invitrogen, NP0007) plus 20 mM Dithiothreitol (DTT, Merck Life Science, 20-265) was added to each sample. Samples were heated for 5 min at 95°C and centrifuged for 1 min at 1,000 × g. Precision Plus Protein Kaleidoscope Standard (BioRad, 1610375EDU) and 15–50 μL of prepared samples were added to the 4–20% Mini-PROTEAN® TGXTM Precast Protein Gels. Proteins were separated for 60–80 min at 120 V. The proteins from the gels were transferred to a nitrocellulose membrane by using Trans-Blot Turbo Transfer System at 25 V for 7 min. After transferring, the membrane was removed and washed in TBST and then blocked in 5% dried milk powder in TBST (binding buffer) at room temperature for 1 h. Membranes were incubated with primary antibody in binding buffer at 4°C overnight. Membranes were washed 3 times with TBST and then incubated with the secondary antibody for 1 h at room temperature in binding buffer. Membranes were washed 3 times in TBST, and then Clarity ECL (Biorad, 170-5061) was added for 1 minute and chemiluminescence was imaged using a Syngene G:Box.

### Imaging of ASC-mCitrine BMDM

Method as previously described by Bertheloot et al^70^. Bone marrow of ASC-mCitrine-expressing transgenic mice was flushed from isolated tibia and femur. Cells were then incubated for 6 days in the presence of L929 culture supernatants generated in-house. Adherent differentiated macrophages were detached from their culture vessel using PBS supplemented with 5 mM EDTA and 2% FBS. Cells were seeded in imaging 96-well plates at 3 × 10^4^ cells per well. Cells were allowed to rest at 37°C, 5% CO_2_ overnight, prior to stimulation. Cells were primed with LPS (100 ng/mL) for 3h and in the presence of k765 (50 µM), then further incubated either at 37°C or 42°C for 1 h. Cells were subsequently stimulated for 2 h at 37°C with nigericin (5 μM), or the combination of the rod protein from Burkholderia pseudomallei fused to lethal factor N-terminus (LFn-BsaK, 100 ng/mL) and Bacillus anthracis protective antigen (PA, 500 ng/mL). Cells were fixed with 4% PFA (in PBS) and nuclei stained with DRAQ5 overnight at 4°C. Finally, cells were imaged on a CellDiscoverer 7 microscope (Zeiss) with 8 positions (2 × 4 images/well) per condition using 6 Z-slices per image. Maximal intensity projections were generated for each image set before the number of cells and specks per field were calculated using Cell Profiler software.

### Imaging of HMDM stained for ASC

Method as previously described by Bertheloot et al^70^. PBMCs were isolated from buffy coats using a density gradient (Ficoll-Paque Plus, GE Healthcare) followed by positive selection of monocytes using a CD14 MACS bead kit (Miltenyi Biotech). Cells were then differentiated into macrophages with 500 U/mL rhM-CSF (Immunotools) for 3 days at 37°C, 5% CO_2_. Adherent cells were harvested using PBS complemented with 2% FBS and 2 mM EDTA. Cells were seeded in imaging 96-well plates at 3 × 10^4^ cells per well. Cells were allowed to rest at 37°C, 5% CO_2_ overnight, prior to stimulation. Cells were primed with LPS (100 ng/mL) for 3 h and in the presence of VX-765 (50 µM), then further incubated either at 37°C or 41°C for 1 h. Cells were subsequently stimulated for 2 h at 37°C with nigericin (5 μM), or the combination of the type III secretion system (T3SS) needle-like protein from Shigella flexneri fused to lethal factor N-terminus (LFn-MxiH, 100 ng/mL) and Bacillus anthracis protective antigen (PA, 1 µg/mL). Cells were fixed with 4% PFA (in PBS) for 30 min at room temperature (RT) and washed twice in PBS. Following permeabilization (in buffer PBS + 2% FBS + 0.5% Triton-X100 + human FcR blocking) for 10 min at RT, cells were stained with PE-labelled anti-ASC antibody (TMS-1, BioLegend, 1 µg/mL) at 4°C overnight in permeabilization buffer. Cells were washed 3-times with permeabilization buffer and nuclei stained with DRAQ5 in PBS for 30 min. Finally, supernatants were exchanged for PBS and cells were imaged on a CellDiscoverer 7 microscope (Zeiss) with 8 positions (2 × 4 images/well) per condition using 6 Z-slices per image. Maximal intensity projections were generated for each image set before the number of cells and specks per field were calculated using Cell Profiler software.

### ASC-speck formation assays

HEK^ASC-EGFP^ expressing NLRP1-HA (HEK^NLRP1+ASC^; cell line H8-1), or NLRP3-HA (HEK^NLRP3+ASC^; cell line H98) under the control of pUbC (kind gifts from Prof. Florian I. Schmidt, Institute of Innate Immunity, Medical Faculty, University of Bonn, Bonn, Germany) were cultured in DMEM containing 10% FBS and 1 × GlutaMax. Cells were seeded in a 24-well plate at 2 × 10^5^ cells per well and allowed to rest at 37°C, 5% CO_2_ overnight. The next day, medium was replaced with 500 μL Opti-MEM and the cells were incubated at either 37°C or 41°C for 3 h. Cells were moved to 37°C and then nigericin (5 μM, 1h) or Talabostat (30 μM, 24h) were added to H98 or H8-1, respectively. The media were removed and 500 μL cold FACS buffer (PBS with 1% FCS and 2 mM EDTA) was added to help detach the cells by pipetting up and down. Cells were filtered through a 100 μm nylon monofilament mesh (Plastok Associates) into a FACS tube. The cells were run through flow cytometer (BD FACS Canto II) using standard operating procedures as previously described^33^. The following parameters were analysed and collected: forward scatter (FSC)-area, FSC-width, side scatter (SSC)-area, FITC-area, FITC -width, FITC -height. The gating and analysis strategies were described as follows: (i) Live cells (FSC-A vs SSC-A); (ii) Single cells (FSC-A vs FSC-W); (iii) ASC specking cells (FITC-A vs FITC-W).

### Culture of HEK293T stably expressing hASC-GFP cells

HEK cells stably expressing hASC-GFP (a kind gift from Prof. Kate Schroder, Institute for Molecular Bioscience, University of Queensland, Australia) were cultivated and maintained in HEK medium: DMEM containing 10% FBS, 1% P/S, 1× GlutaMax and blasticidin (4 μg/mL). Cells were seeded in 24-well plates at 2 × 10^5^ cells in 500 μL HEK medium (without blasticidin) per well and allowed to rest at 37°C, 5% CO2. 5 h after cell seeding 300 μL of media was removed from each well. Cells were then transfected with 100 ng of empty vector plasmid or mCherry tagged NLRP3/mutants using 0.3 μL Lipofectamine 2000 per well. The plate was centrifuged at 1000 × g for 10 min at room temperature to increase transfection efficiency. The transfected cells were incubated at 37 °C with 5% CO2 for 16 h.

### Measurement of ASC speck formation by Dual-Colour TOFIE

16 h after transfection, medium was removed and 300 μL of Opti-MEM was added, followed by the addition of R837 (100 μM, 4 h) or Nigericin (5 μM, 1 h). The media were removed and 500 μL cold FACS buffer was added to help detach the cells by pipetting up and down. Cells were filtered through a 100 μm nylon mesh into a FACS tube. The cells were analysed by flow cytometry (BD FACSAria™ III Cell Sorter) using standard operating procedures with the following parameters being analysed and collected: forward scatter (FSC)-area, FSC-width, side scatter (SSC)-area, FITC-area, FITC-width, FITC-height, mCherry-area, mCherry-width and mCherry-height. The gating and analysis strategies were as follows: (i) Live cells (FSC-A vs SSC-A); (ii) Single cells (FSC-A vs FSC-W); (iii) GFP+ mCherry+ (FITC-A vs mCherry-A); (iv) mCherry expression subgates (‘Very Low’, ‘low’, ‘medium’ and ‘high’ gates in Q2); (v) ASC specking cells (FITC-A vs FITC-W). The subpopulation of very low mCherry expression in (iv) were used for ASC speck quantification by the time-of flight inflammasome evaluation (TOFIE) method as previously described^40^

### Cellular thermal shift assay (CETSA)

BMDM were seeded at 3-5 × 10^5^/mL, 10 mL in 10 cm dishes. The next day cells were pretreated with a caspase-1 inhibitor (25 μM Ac-YVAD-cmk or 20 μM VX-765) and were stimulated with LPS (100 ng/mL) 1 h, or LPS and nigericin (5 μM) simultaneously for 1 h, all at 37°C. HEK293T cells were seeded at 3 × 10^5^/mL in 10 cm dishes, the next day they were transfected with 3 μg of plasmid DNA for NLRP3-Twin-Strep-tag (TST) or NLRP3Δ177-195-TST using the calcium phosphate method^71^. CETSA methods are based on Jafari et al^72^. Cells were harvested by scraping and rinsed in PBS. Cells were resuspended in PBS containing protease inhibitors (Proteoloc Protease Inhibitor Cocktail EDTA-free, Abcam, ab270055) and transferred to thin walled PCR tubes on ice. CETSA was performed by heating cells between 40-64°C in a Veriti 96-well Thermocycler (Applied Biosystems). Cells were lysed by addition of 6% Triton X-100 solution on ice for 20 min with repeated vortexing. Samples were then centrifuged at 20,000 × g for 10 min at 4°C. Clarified lysate was removed and mixed with 4 × LDS loading buffer and 20 mM DTT. Samples were heated at 95°C for 5 min and briefly spun down before being analysed by immunoblotting.

### Plasmid preparation

The pEF6 mammalian expression vector with human NLRP3 mCherry was a kind gift from Dr Kate Schroder^40^. Point mutations were generated using a Q5® Site-Directed Mutagenesis Kit (New England Biolabs). Sanger sequencing of plasmids was performed at Eurofins Genomics. Plasmid synthesis was performed at Genscript for pcDNA3.4 vectors containing human NLRP3 with a C-terminal twin-Strep-tag or mCherry tag. In the NLRP12 swap mutant NLRP3 amino acids 172-207 (RLIKEHRSQQEREQELLAIGKTKTCESPVSPIKMEL) are replaced with NLRP12 amino acids 161-198 (LLVKEHSNPMQVQQQLLDTGRGHARTVGHQASPIKIET). Plasmids were prepared for transfection with a PureLink HiPure Plasmid Maxiprep Kit (Invitrogen). Primers were synthesised by Integrated DNA Technologies.

### Molecular modelling and structural bioinformatics

Molecular dynamics (MD) simulations were carried out using the following experimental structures: human NLRP3 in closed/autoinhibited conformation (PDB code: 7PZC, chain A), human NLRP3 in active/open conformation (PDB code: 8EJ4). Complexes were visually inspected, and structures were prepared using UCSF Chimera^73^, by removing small molecules (water molecules, ions, inhibitors and cofactors bound), extracting a single monomer, removing N-terminal PYD and linker (residues 1-134), and modelling missing side chains. Missing loops were modelled using AlphaFold3^74^ and fitted using UCSF Chimera^73^. Initial structure of G191A and τιCOFI mutants of human NLRP3 NACHT domain have been modelled in UCSF Chimera^73^. Residues 177-195 were modelled separately using AlphaFold3^74^, swapped in closed conformations, and the backbone has been linked. In parallel, all investigated variants of NLRP3 (WT, G191A, τιCOFI and NLRP12-swap) were modelled using AlphaFold3^74^. All structures have been subjected to up to 50,000 steps of molecular-mechanical energy minimisation in Gromacs^75^, followed by 100 ns all-atom MD simulations in triplicate, at standard temperature (300 K). Highest-populated clusters from open and closed conformations were selected as inputs for BEE/eBDIMS workflow. For NLRP3 closed and open conformations, clusters were used to follow up with coarse-grain MD at higher temperatures. Structure-based predictions of aggregation properties were carried out using Aggrescan3D (A3D v.2.0), which considers fluctuations in the protein’s intrinsic dynamic fluctuations^76^. Primary sequence alignments were performed using ClustalW online tool^77^.

### Coarse-grained molecular dynamics

The NLRP3 constructs (monomers lacking PYD; residues 1-134 removed) in either closed (inhibited) or open (active) conformations of NACHT domain were converted to coarse-grained (CG) representation using the *martinize2*.*py* workflow module^78^ of the MARTINI 3 force field^79^. The elastic network model was applied to reinforce the stability of the secondary structure of NACHT domain (structured regions) and LRR domain. Default values of the force constant of 500 kJ/mol/nm^2^ were used, with the lower and upper elastic bond cut-off to 0.5 and 0.9 nm, respectively. CG simulations were performed using Gromacs^75^. The pH of the systems was considered neutral. All the simulations were run in presence of regular MARTINI water, were neutralised and brought up to 0.1 M NaCl. The structures were equilibrated for 500 ps. The long-range electrostatic interactions were used with a reaction type field having a cutoff value of 1.2 nm. We used potential-shift-Verlet for the Lennard-Jones interactions with a value of 1.2 nm for the cutoff scheme and the V-rescale thermostat with reference temperatures of 300 K, 360 K, and 400 K in combination with a Berendsen barostat with a coupling constant of 1.0 ps, compressibility of 3.0 × 10^−4^ bar ^−1^, and a reference pressure of 1 bar was used. The integration time-step was 20 fs. All the simulations were run in triplicate for 1 μs. For further analysis and comparison, the extracted CG structures were then converted to all atomistic (AA) representation using in-house script and CG2AT2 tool^80^. The RMSD, RMSF, and radii of gyration were calculated using Gromacs toolkit^75^. The trajectories were followed up with 100 ns all-atom MD simulations (Gromacs) of the NACHT domain at the target temperature.

### Brownian Dynamics – Enriched Ensembles (BEE)

BEE is an in-house multiscale workflow developed to assess large conformational changes of large multiprotein complexes. It is designed to predict protein intermediate conformational states, “invisible” to experimental techniques, from structurally rich ensembles using principal component analysis (PCA), utilising hybrid elastic-network Brownian dynamics and all-atom MD simulations. Firstly, BEE employed the in-house version of Elastic Network Driven Brownian Dynamics Importance Sampling (eBDIMS)^81^ code to perform coarse-grained simulations on two distinct conformers of the target (protein of interest, such as NACHT domain of NLRP3). This process enabled the generation of a transition pathway between these two conformers. PCA or Normal Mode Analysis (NMA) were applied to obtain the modes and frequencies of protein motions: while PCA selected the two principal components with the largest eigenvalues as the pathway for conformational transformation; NMA selected the lowest-frequency motions as the transition pathway. Both approaches are effective for validating transition pathways in terms of conformational changes. Subsequently, PD2^82^ and Scwrl4^83^ were invoked to perform reverse mapping on the whole coarse-grained trajectory, resulting in the generation of protein structures at the full atomic level. These structures were then subjected to the refinement by all-atom MD simulations in GROMACS^75^. For NACHT domain of human NLRP3, 20 intermediates were generated for each conformational transition (closed-to-open and open-to-closed, respectively), which were used to generate inputs for simulations of isolated COFI regions.

### Atomistic molecular dynamics simulations

All simulations used Gromacs^75^ with AMBER99SB-ILDN^84^ parameters. Each simulated COFI region was immersed in a cubic TIP3P water box, set to be 1 nm away from the edge of the protein. 3D-PBC were applied. Na^+^ and Cl^−^ ions were added to maintain the charge neutrality of simulated units. Prior MD simulations, each system underwent energy minimisation via the steepest descent method for 1,000 cycles and the conjugate gradient method for further refinement, with the energy step size of 0.001 nm and a maximum of 50,000 steps. The minimisation was concluded when the maximal force descended below 1000 kJ/mol/nm. Long-range electrostatic interactions were addressed using the Particle-Mesh Ewald (PME) method^85^, while a cut-off of 1.0 nm was applied for short-range electrostatic and van der Waals interactions. Following energy minimisation, all systems were subjected to a 500 ps NVT equilibration with a step size of 2 fs. The protein and non-protein groups were gradually heated to the target temperature (300, 360 and 400 K, respectively) under the influence of the V-rescale thermostat with a time constant of 0.1 ps. LINCS (Linear Constraint Solver) position restraints were applied to the bond lengths and angles of the backbone atoms. The non-bonded short-range interactions were treated with the Verlet cut-off scheme, setting the cut-off distance to 1.0 nm. Long-range electrostatics were once again addressed with PME. Subsequently, NPT equilibration was performed, where temperature was maintained at the target with the continued utilisation of the temperature coupler, followed by the initiation of Parrinello-Rahman pressure coupling^86^ for 500 ps of pressure equilibration, with the target pressure established at 1 bar. Post-equilibration, the systems were subjected to three separate 100 ns simulations to obtain trajectories for analysis, discarding the initial 10 ns of data during the final evaluation. Trajectory analyses were conducted using Gromacs tools. The overall stability was evaluated by RMSD, and local flexibility was assessed via per-residue RMSF. PCA was employed to explore the key motion modes. Secondary structure changes were quantified by DSSP, visualised using a custom in-house script. The protein overall structural stability was analysed through the radius of gyration (Rg). Examination of hydrogen-bonding network was carried out utilising UCSF Chimera^74^ and VMD^87^. Protein-protein interaction enthalpy calculations were performed using parameters derived from AMBER parm99 classical molecular mechanical force fields and a GB/SA implicit solvation model. All calculations were performed using INTAA webserver^88^. For each temperature, trajectories were subjected to cluster analysis and the highest-populated clusters were used as inputs for well-tempered metadynamics.

### Well-tempered metadynamics

Well-tempered metadynamics simulations were carried out using PLUMED 2.90^89^. For simulations of COFI regions, end-to-end distance were defined as collective variable. The initial height of each Gaussian potential (HEIGHT) was set to 0.6 kJ/mol, with a width (SIGMA) of 0.1 nm, and a Gaussian added every 200 simulation steps (PACE). A bias factor (BIASFACTOR) of 10 was established to realise a well-tempered sampling strategy. The addition of Gaussians was based on the current value of the collective variable and was recorded in the HILLS file, to facilitate subsequent analysis and reproduction of the simulation process. Free energy profiles were obtained by integrating data collected during the metadynamics simulations. Comparing the free energy minima and peaks of each allows comparison of interactions stabilising COFI regions at different temperatures and linking those to observed conformations of the NACHT domains and activation states of NLRP3.

### Mouse heating and LPS challenge model

Twelve-week old male C57BL/6 mice were obtained from Australian Bioresources (ABR). Mice were heated as described in Munoz et al^43^. Briefly, the animals were housed at 38°C in a custom-made chamber for 18 h or at 22°C (control room temperature). Core body temperature was measured immediately before and after heat-treatment using a rectal probe, and heated mice were allowed to recover for 2 h at 22°C before LPS challenge. Mice were injected intraperitoneally with 100 μg of LPS (from *Escherichia coli* O111:B4, Sigma) and culled 2 h later for the collection of blood by retro-orbital bleeding. Serum was prepared by incubating the blood at room temperature for 20 min to allow coagulation before centrifuging at 10,000 × g for 15 min at 4°C to remove the clot. The serum was snap-frozen in liquid nitrogen and stored at −80°C for later analysis. Inflammatory cytokines and chemokines in serum were measured using the Bio-Plex Pro Mouse Cytokine 23-plex Assay (BioRad, M60009RDPD) with a LuminexMagPix® cytoplex platform according to the manufacturers’ instructions. IL-18 was measured using an ELISA for mouse IL-18 (Abcam, ab216165).

### Key resources table

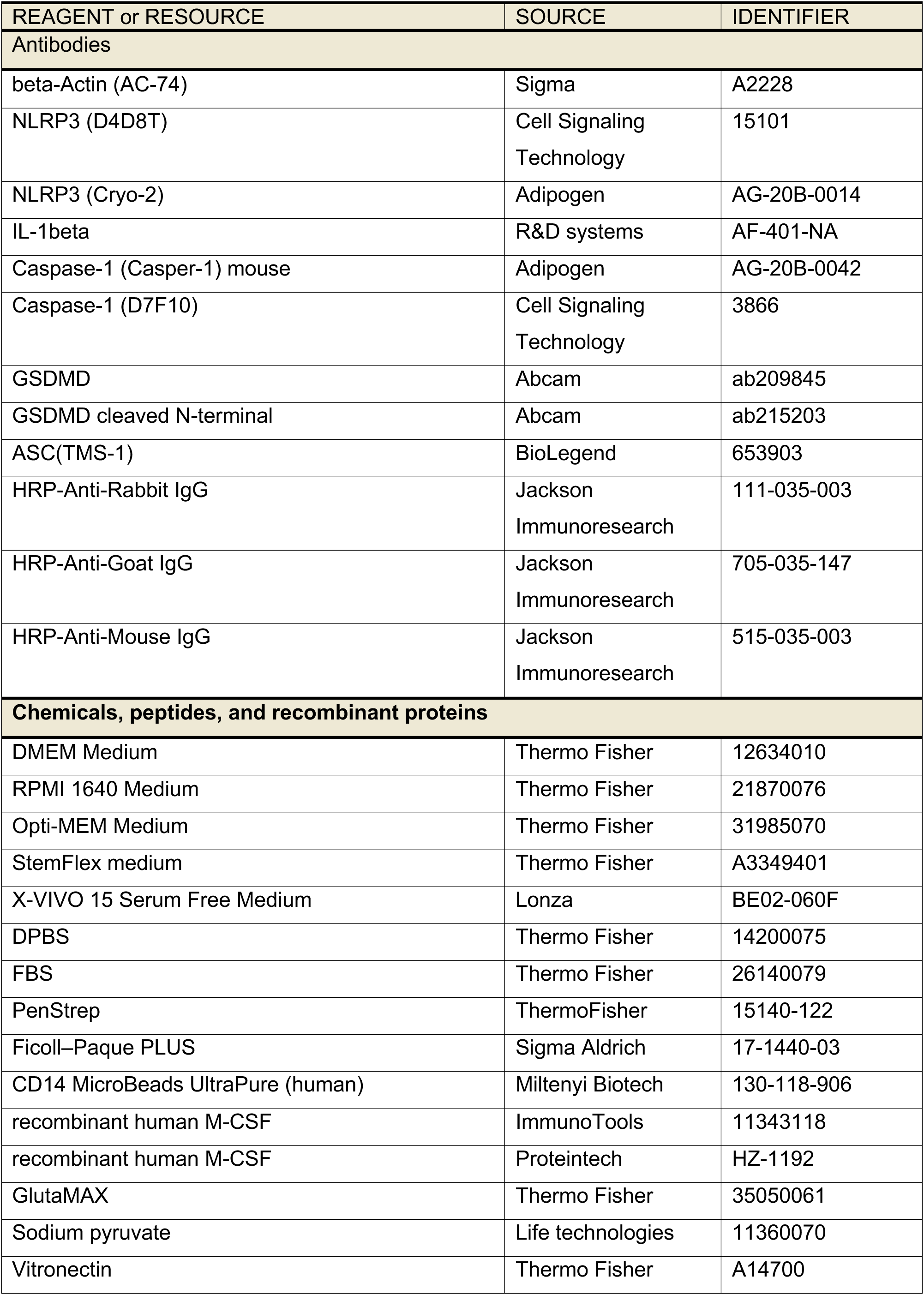
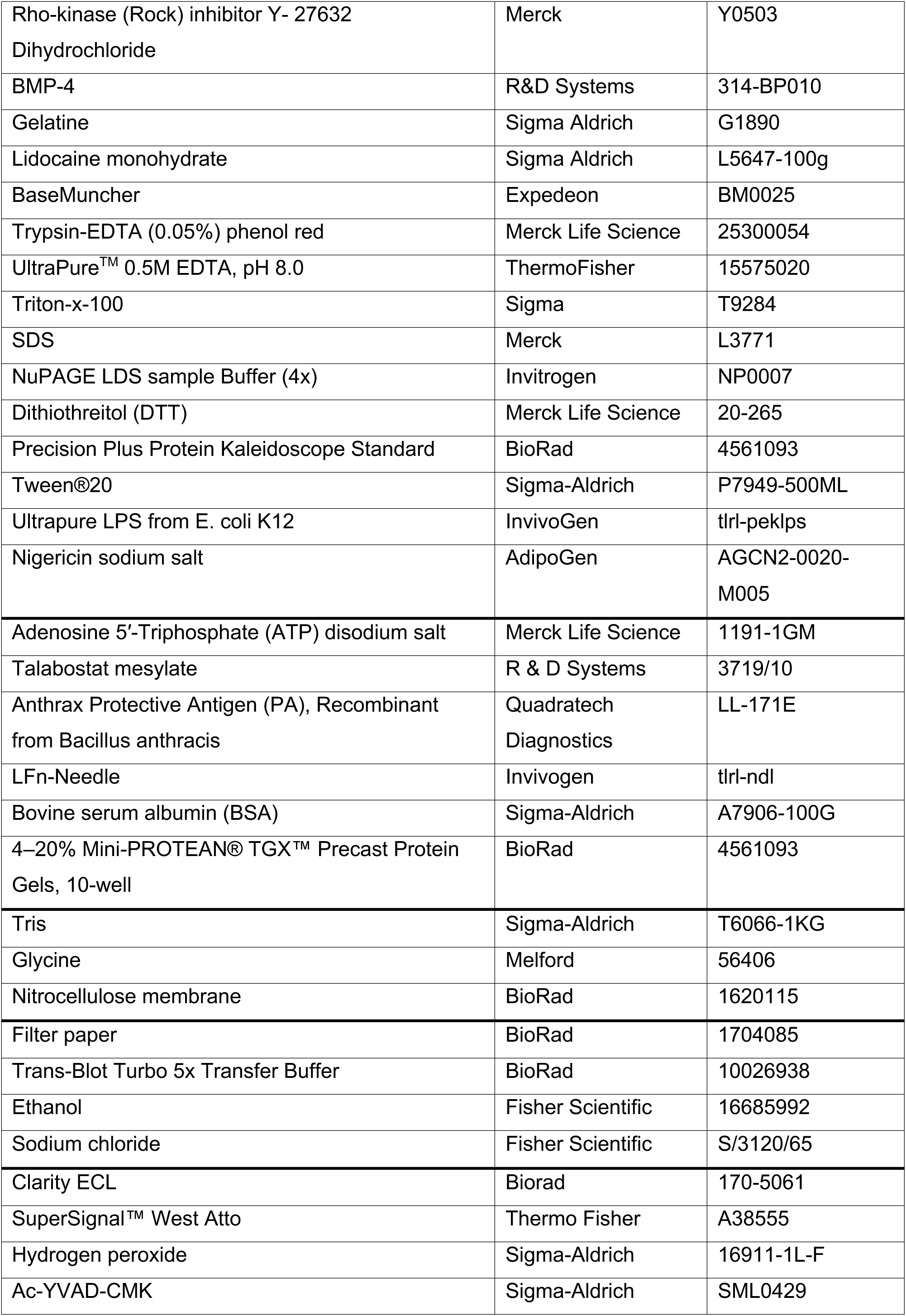
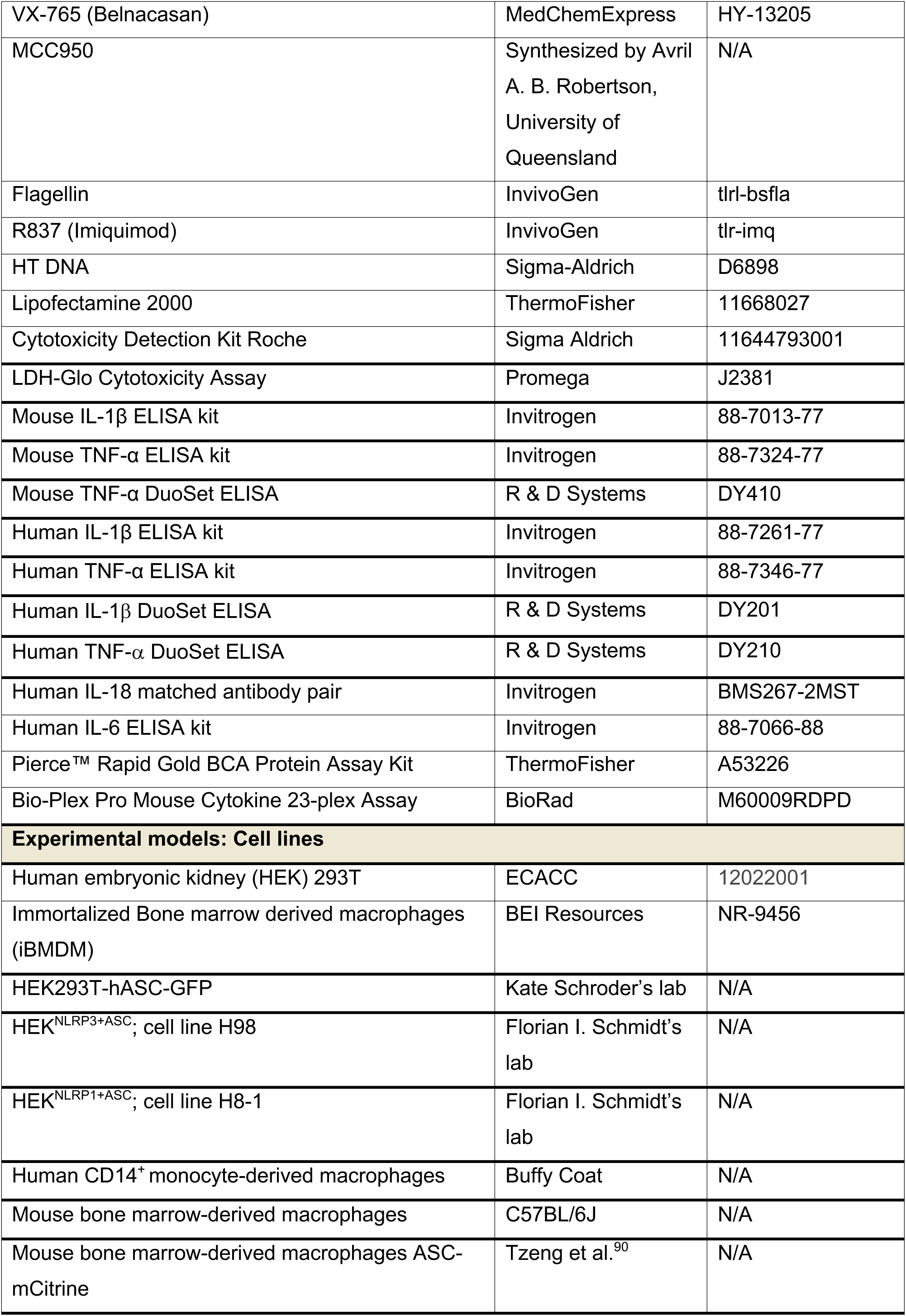
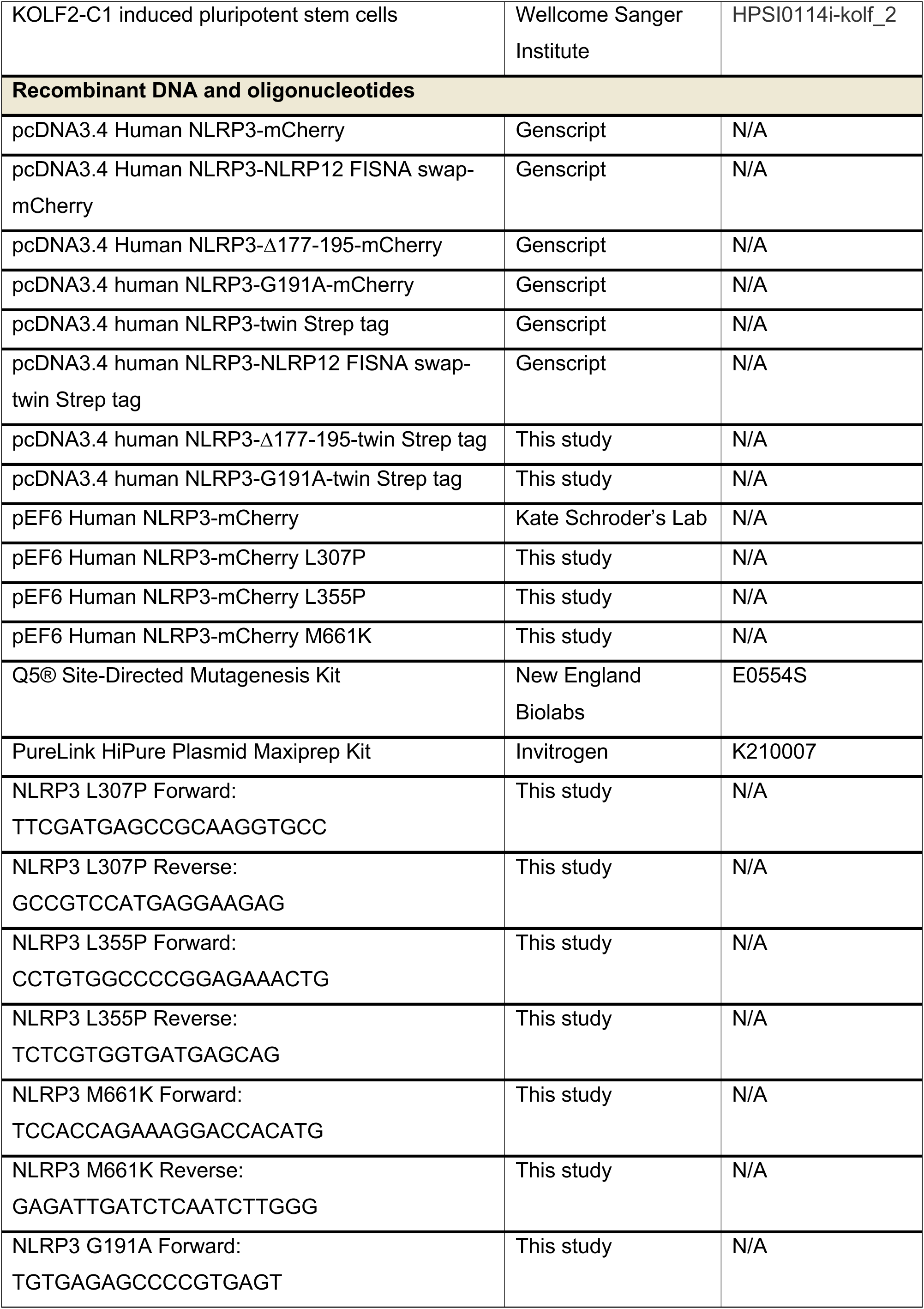
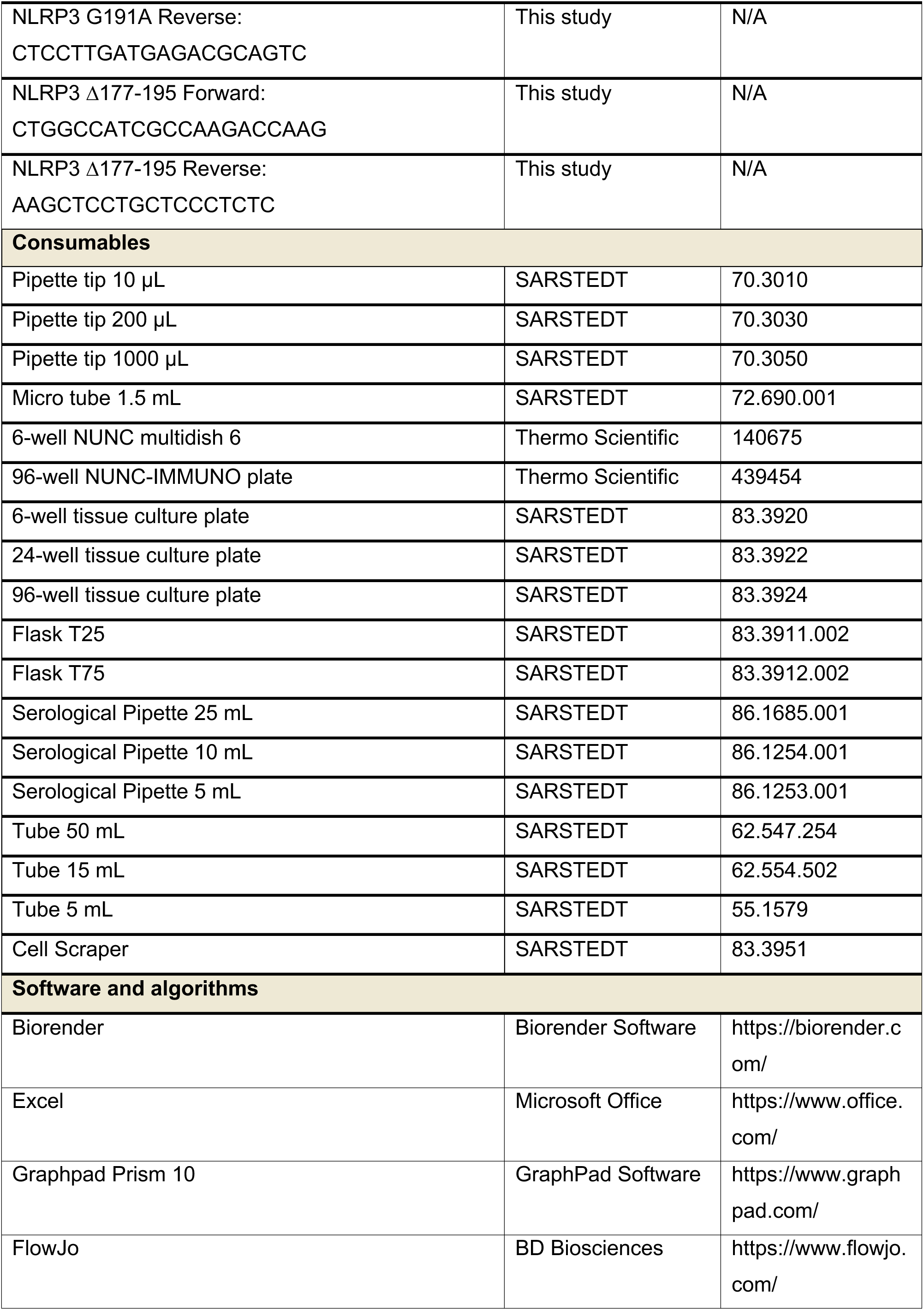
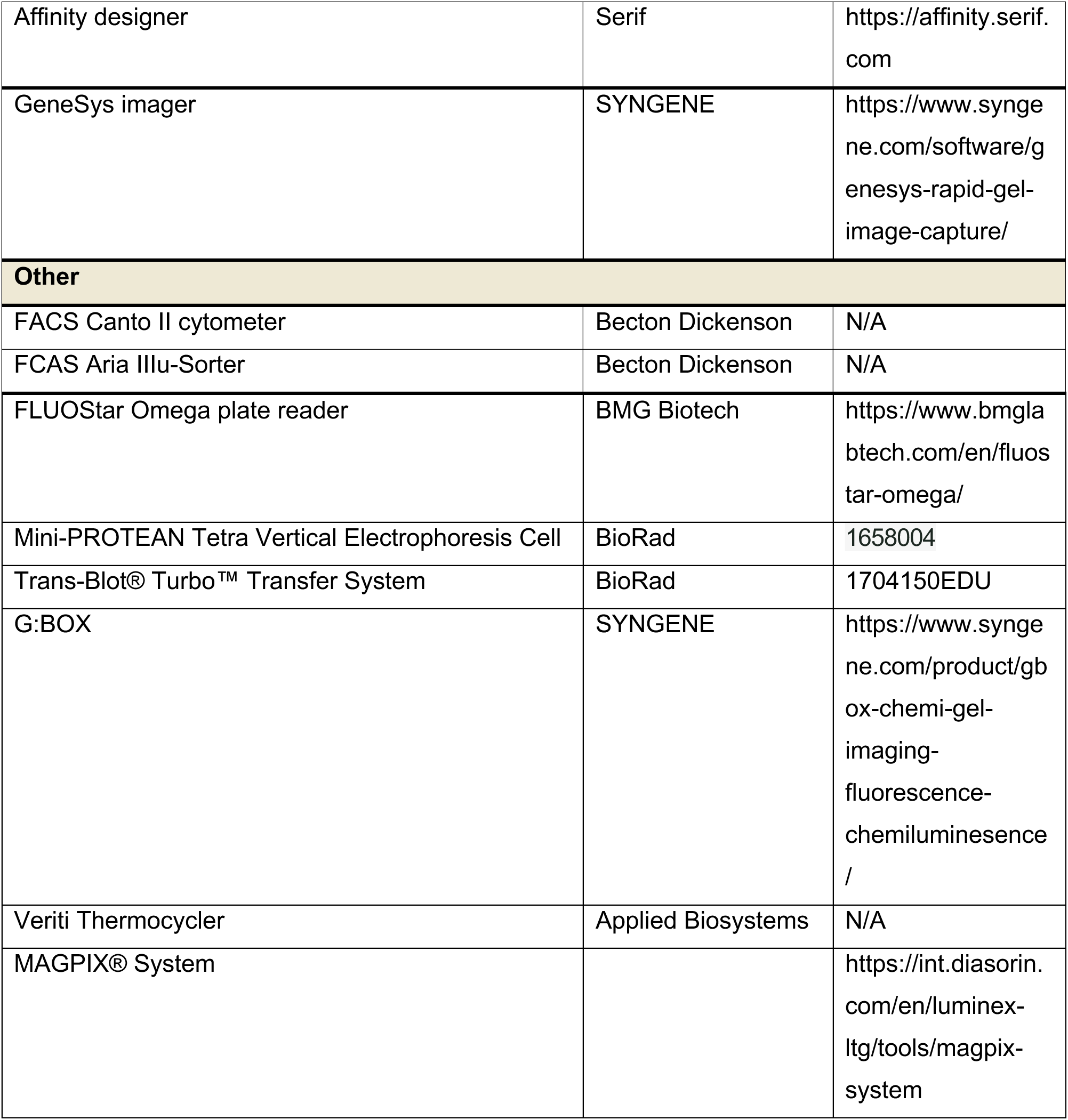

